# Specificity and exon target space of splicing modifying compounds

**DOI:** 10.1101/2025.08.14.670273

**Authors:** Felina Lenkeit, Judith Knehr, Marc Altorfer, Andrea Byrnes, Wenjing Li, Jack Hsiao, Connie Wu, Priti Gaitonde, Philip R. Skaanderup, Steve Mullin, Elizaveta Solovyeva, Michal Pikusa, Andrew T. Krueger, Johannes Ottl, Caroline Gubser Keller, Christian Kolter, Ulrike Naumann, Philipp Ottis, Alejandro Reyes

## Abstract

Modulation of splicing has become an established therapeutic strategy, with proven clinical applications and continued potential to target specific exons to influence gene expression. Recent advancements led to the identification of small molecule splicing modifiers such as Risdiplam and Branaplam. These compounds induce the inclusion of exons that are typically skipped due to their weak 5′ splice site. While Risdiplam has a preference to induce exons with a N_−3_G_−2_A_−1_ sequence at the 3′ exon end, Branaplam has a proclivity towards introducing A_−3_G_−2_A_−1_ -ending exons. However, the variables that determine the selectivity and specificity of splicing modulators are still not completely understood, as evidenced by the hundreds of unaffected N_−3_G_−2_A_−1_ -ending exons present in the human genome. In this study, we delve into the molecular mechanisms governing the specificity of splicing-modifying compounds, focusing on their interactions with RNA structures at splice sites. Using biochemical assays, whole transcriptome analyses, and genetic perturbation approaches, our findings reveal contributions of primary sequence codependencies that help determine the required secondary structural conformation, thus governing responsiveness to splicing modulator induction. Based on these learnings, we were able to reprogram the specificity of splicing modulators by genetic manipulation of the U1 snRNA component of the spliceosome. Our findings further the understanding of splicing modulators and might help to identify new targets and compounds.

## Introduction

Splicing is a fundamental biological process that generates mature mRNA by removing introns from premRNA. Splicing is carried out by the spliceosome, a macromolecular complex which comprises various protein components and small nuclear RNAs (snRNAs), together termed small ribonucleoproteins (snRNPs). One of these snRNAs, as part of the U1 snRNP, guides the first step of the splicing reaction by base-pairing with the sequence spanning from the terminal three nucleotides at the 3*′* end of an exon (positions -3, -2, -1) to the eight nucleotides of the downstream intron (positions +1 to +8) (Wilkinson et al., 2020; Papasaikas and Valcarcel, 2016) (Figure 1A). This sequence stretch in the premRNAs is known as U1 binding site and its consensus sequence is 5*′*-C_−3_A_−2_G_−1_G_1_U_2_A_3_A_4_G_5_U_6_A_7_U_8_-3*′*. Similarly, the branchpoint sequence, the polypyrimidine tract and the 3*′* splice site (3*′*ss) are sequence elements that guide the splicing machinery to determine the 5*′* start of an exon. In most human genes, alternative selection of 5*′* and 3*′* splice sites enable the production of multiple isoforms from a single gene, contributing to protein diversity (Black, 2003; Lee and Rio, 2015).

**Figure 1.**
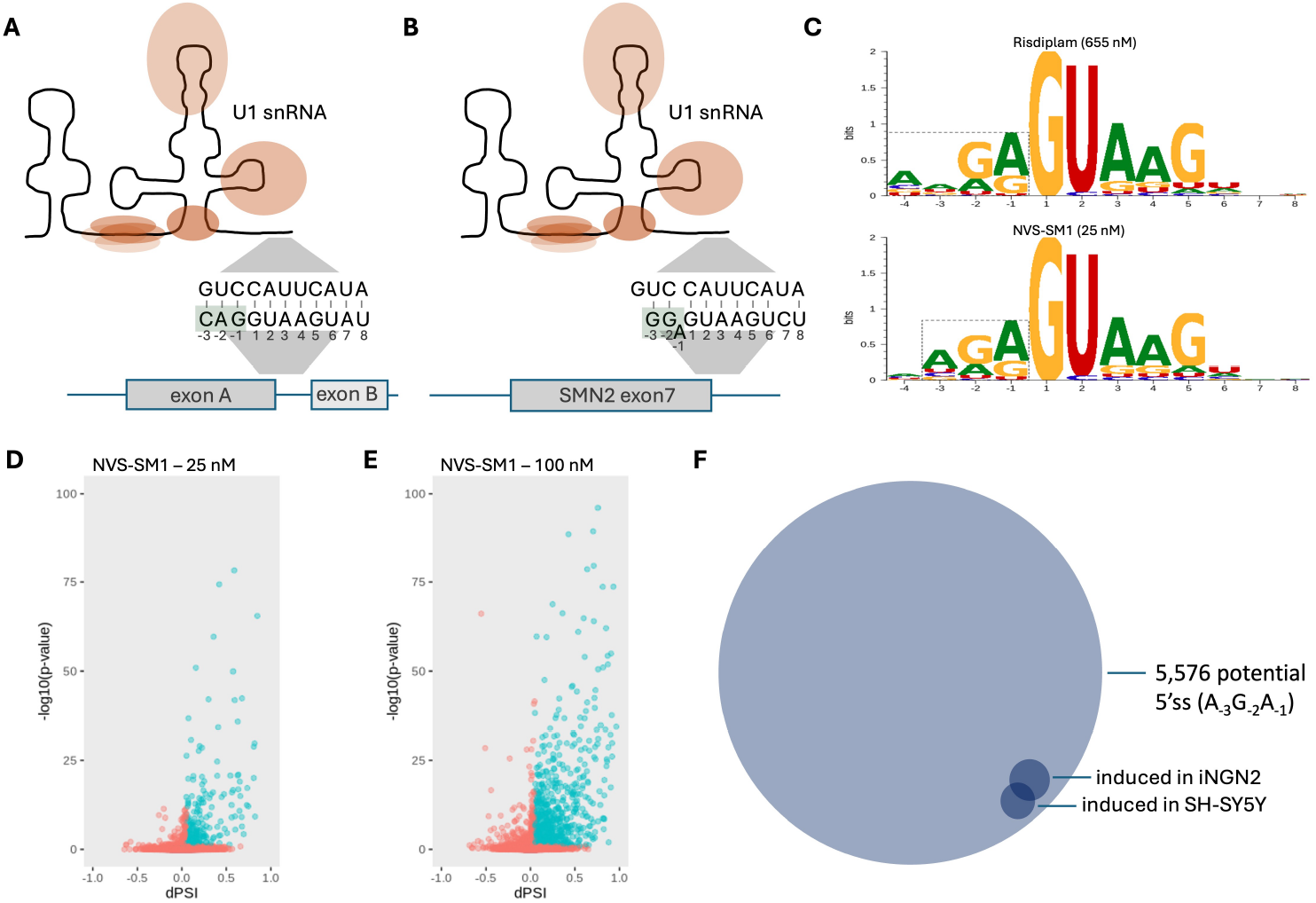
Sequence specificity of splicing compounds. (A) Splice site recognition by base-pairing between the U1 snRNA within the spliceosome and an exon carrying a consensus U1 binding site (here called exon A). The U1 binding site spans from the three nucleotides at the 3^*′*^ end of exon (positions -3, -2, -1) to the eight nucleotides of the downstream intron (positions +1 to +8). (B) Splice site recognition of exon 7 in the *SMN2* gene in the presence of a splicing modifier like Risdiplam. (C) RNA sequence logos to represent sequence conservation at the U1 binding site of all exons included in response to either 655nM Risdiplam, or 25nM NVS-SM1. The last 4 nucleotides of the exon (position -4 to -1) and the first 8 nucleotides of the following intron (position +1 to +8) are included in the logo. (D) Differential splicing analysis between whole transcriptome profiles of SH-SY5Y cells treated with 25 nM NVS-SM1 and SH-SY5Y cells treated with DMSO. Exons highlighted in turquoise are considered as included in response to compound treatment (dPSI > 0.05, adjusted p-value < 0.1) (E) Differential splicing analysis between whole transcriptome profiles of SH-SY5Y cells treated with 100 nM NVS-SM1 and SH-SY5Y cells treated with DMSO. (F) Venn diagram to highlight gap between 5,576 potential 5^*′*^ss matching primary sequence preference of NVS-SM1 and the actual number of detected induced AGA-ending exons in response to NVS-SM1 treatment in SH-SY5Y and iNGN2 cells.

Modification of splice site selection has been proven to be a promising approach in treating various diseases. This strategy addresses not only diseases caused by mutations that lead to aberrant splicing, but also diseases with other causes, where modulating the splicing process can influence the expression levels of particular isoforms (Tang et al., 2021). By directly targeting mRNAs, this approach enables the modulation of proteins that may otherwise be challenging to drug or even considered undruggable. While antisense oligonucleotides (ASOs) constitute the majority of clinically validated splicing modifiers (Mercuri et al., 2018; Havens and Hastings, 2016; Mendell et al., 2013), RNA-targeting small molecules offer numerous advantages. Notably, these small molecules allow non-invasive oral delivery, ensure systemic distribution, and have the capacity to cross the blood–brain barrier, thus enhancing their potential as therapeutic agents (Warner et al., 2018; Singh et al., 2020).

To date, there are several small molecule splicing modulator compounds with available human clinical data, including Risdiplam, NVS-SM1 (Branaplam) (Borowsky et al., 2026), PTC518, SKY-0515, RGT-61159 and REM-422. Risdiplam is being used for the treatment of spinal muscular atrophy (SMA) (Dhillon, 2020). SMA is caused by a mutation in the survival motor neuron 1 (*SMN1*) gene that leads to a deficiency of functional SMN proteins (Lefebvre et al., 1995). The gene *SMN2* is nearly identical to *SMN1*, however harbors a critical C to T substitution that deviates the 5*′*ss of exon 7 from the canonical sequence. This weak 5*′*ss (5*′*-G_−3_G_−2_A_−1_G_1_U_2_A_3_A_4_G_5_U_6_C_7_U_8_-3*′*) prevents efficient splicing of exon 7 and as a result, most *SMN2* produce non-functional isoforms (Figure 1B). By introducing exon 7 into transcripts of *SMN2*, Risdiplam restores the expression of functional SMN protein that compensates for the *SMN1* pathologic mutations (Ruggiu et al., 2012; Le et al., 2005; Ratni et al., 2018). Branaplam, here referred to as its development code NVS-SM1, was also found to promote exon 7 inclusion in *SMN2* transcripts. Additionally, this molecule demonstrated the capability to induce the inclusion of a frame-shifting pseudoexon between exon 49 and 50 in the *HTT* gene that introduces a pre-mature stop codon which triggers the mRNA to be degraded (Bhattacharyya et al., 2021; Keller et al., 2022; Chen et al., 2020). This *HTT* pseudoexon is not recognized by the spliceosome in the absence of NVS-SM1, as it contains a non-canonical 5*′*ss (5*′*-A−3G−2A−1G1U2A3A4G5G6G7G8-3*′*).

Genome-wide transcriptome analyses and mutational screening assays have shown that Risdiplam has a preference to induce exons with a N_−3_G_−2_A_−1_ (N = any base) sequence at the ^*′*^ exon end, while NVS-SM1 has a proclivity towards introducing A_−3_G_−2_A_−1_- ending exons (Krach et al., 2022; Ishigami et al., 2024) (Figure 1C). Without the presence of these compounds, these exons are typically being skipped due to their weak 5*′*ss, making their recognition unlikely (Monani et al., 1999; Wong et al., 2018; Freund et al., 2005). An NMR structure of a chemical analog of Risdiplam, SMN-C5, showed that the compound binds the RNA helix formed by the 5*′*ss and the 5*′*end of the U1 snRNA in its major groove. Here, it stabilizes the bulged-out A_−1_ at the exon end, acting like a molecular glue that strengthens the interaction of both RNAs, compensating for the weak 5*′*ss to enable splicing initiation (Campagne et al., 2019; Ratni et al., 2016). Subsequent NMR analyses extended this bulge-repair mechanism to diverse chemotypes, including Branaplam analogs, confirming that structurally distinct compounds can target the same A_−1_ bulge at the exon-intron junction (Malard et al., 2024). Beyond structural models, Ishigami and co-workers applied a quantitative high-throughput splicing assay and modeling approach to demonstrate that, while Risdiplam recognizes a relatively narrow N_−3_G_−2_A_−1_ motif, Branaplam displays two distinct sequence-recognition modes at the ^*′*^ss. This suggests that specificity arises from the subtle interplay between local sequence composition and RNA conform2ational dynamics (Ishigami et al., 2024). Moreover, they identified the –4 position as a key determinant within the A_−4_N_−3_G_−2_A_−1_ motif of Risdiplamresponsive exons. With all analyses performed in a fixed genetic background, where exon architecture, U1 snRNP composition and spliceosomal components remained unaltered, the influence of these characteristics in different local genetic contexts is still unexplored.

While the studies above have substantially contributed to our understanding of small-molecule splicing modulation, the determinants of the underlying selectivity remain incompletely resolved. For instance, a recent study predicted >58,000 potential exons with a 5*′*ss sequence of 5*′*-A_−1_G_−2_A_−3_G_1_U_2_A_3_A_4_G_5_-3*′* in human introns, however observed inclusion upon NVS-SM1 treatment for only 609 exons (Bhattacharyya et al., 2021). Similarly, transcriptome data have shown that N_−3_G_−2_A_−1_ ending exons are induced at different levels upon treatment with NVS-SM1 or Risdiplam.

Here, we analyzed the contribution of sequence elements to the specificity of splicing modulators using a combination of biochemical assays, whole transcriptome analyses, and genetic perturbation approaches. In contrast to prior studies focused primarily on the structural mechanism of bulge stabilization, we provide quantitative evidence linking sequence complementarity between U1 snRNA and the exon to drug responsiveness. Beyond characterizing exons that are induced by splicing modulators, we also investigated exons that remain unresponsive to treatment in order to reveal sequence features that prevent compound activity. Our analyses suggest that the presence of an inducible A_−1_ bulge is not merely a structural consequence of compound binding but rather a prerequisite for drug-mediated exon inclusion. To address additional determinants of specificity, we examined whether the conservation of nucleotides in the intronic portion of U1-binding sites among compound-responsive exons arises from direct involvement in compound–RNA interactions or instead reflects their role in stabilizing the U1–5*′*ss duplex duplex. By genetically modifying the U1 snRNP, we demonstrate that sequence complementarity between U1 snRNA and target exons in these intronic nucleotides not only drives sensitivity but can also be exploited to modulate compound specificity. Finally, we systematically integrated sequence context and predicted regulatory elements and developed a machine learning framework to evaluate their contribution to compound responsiveness.

Taken together, these results refine the parameters that define the splicing modulator target space and provide mechanistic insights for the rational design of nextgeneration splicing-modifying compounds.

## Results

NVS-SM1 is known to preferentially induce the splicing of exons with 3*′* ends consisting of an A_−3_G_−2_A_−1_ trinucleotide (Figure 1C), which we refer to as AGA exons for simplicity (Keller et al., 2022). First, we aimed to define the potential space of AGA target exons throughout the human genome. We used the large collection of human RNA-seq data from the Genotype-Tissue Expression Project available through the snaptron database (Wilks et al., 2018). We identified 5,576 potential 5*′*ss matching a 5*′*-A_−3_G_−2_A_−1_G_1_U_2_-3*′* motif. Most of the corresponding exons were supported by unannotated exon-exon junctions and, when detected, were included at low level, confirming that these non-canonical exons are usually not recognized by the splicing machinery under normal physiological conditions.

### Variation of exon target space of NVS-SM1 across cell types

This large number of predicted target exons, however, is in strong contrast to the reported transcriptomic profiles in literature, where a few hundred of AGA exons are reported to be induced in response to splicing modifiers such as NVS-SM1 (Keller et al., 2022; Krach et al., 2022). Consistent with these reports, we detected the induction of 125 and 304 AGA-ending exons when we treated SH-SY5Y cells for 24 hours with 25nM and 100nM of NVS-SM1, respectively, in whole transcriptome profiles (10% false discovery rate [FDR]; Figure 1D/E). Despite focusing on non-canonical 5*′*ss, we detected 334 AGA exons with high levels of baseline inclusion in the absence of a compound (>0.6 percent spliced-in [PSI], Supplementary Figure S2A). Notably, the induction of these high baseline AGA exons was not further boosted upon compound treatment.

To further refine the AGA exon target space, we sequenced the transcriptomes of SH-SY5Y, as well as iNGN2 cells, in a single experiment, following treatment with 25nM of NVS-SM1 for 24 hours. Compared to DMSO controls, we found 97 induced AGA exons in SHSY5Y cells and 171 in iNGN2 cells, with 75 of these exons common to both (Figure 1F). When focusing on the list of genes containing AGA-ending exons induced only in SH-SY5Y cells, these genes, on average, demonstrated a 90-times higher expression in SH-SY5Y, compared to iNGN2 cells, and vice versa (Supplementary Figure S2B). These observations suggest that a large proportion of AGA exons detected in only one cell type are explained by cell type specific expression of genes containing induceable AGA exons. Indeed, when we filtered the 5,576 potential AGA exons for those harbored by highly expressed genes in SH-SY5Y cells and, at the same time, demonstrating low baseline inclusion in the absence compound, the number of potentially inducible AGA exons in SY-SY5Y cells dropped to 628.

Even when considering AGA exons with low baseline exon inclusion and high gene expression (where detectability is not an issue due to low gene expression), approximately half of the potentially inducible AGA exons remained uninduced following NVS-SM1 treatment. We reasoned that if the induction of AGA exons introduced stop codons into transcripts, a rapid degradation of mRNA by nonsense mediated decay (NMD) could prevent their detection by RNA-seq. However, only 8.5% of the genes carrying an uninduced AGA exon were downregulated (Figure 2A), indicating that this effect would explain only a minority of the unobserved AGA exons. To confirm this observation, we knocked down major RNA decay pathways and analyzed the splicing patterns upon compound treatment. Compared to compound treatment alone, the knocked down cells showed only a tiny increase in AGA exon inclusion (Supplementary Figure S2C, mean delta PSI 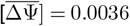), supporting the notion that RNA degradation does not explain the majority of unobserved AGA exons.

**Figure 2.**
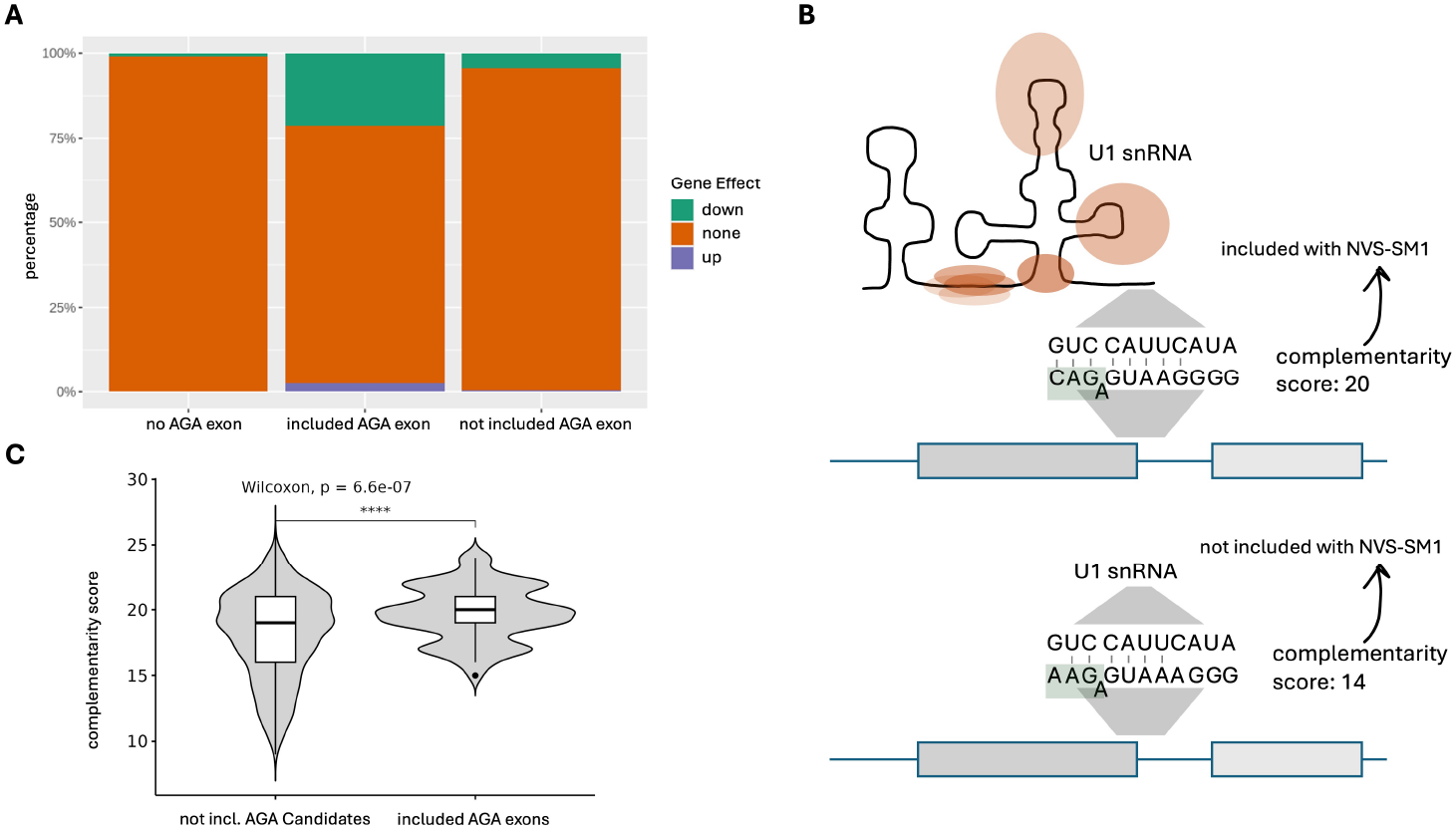
Compound effects on gene expression and U1 snRNA complementarity in exon inclusion. (A) Effect on gene expression (upregulation, downregulation or no effect) of NVS-SM1 treatment of SH-SY5Y cells. 8.5% of the genes carrying an AGA exon, not inducible upon NVS-SM1, are downregulated. (B) Example of two AGA exons that seem to be good candidates for NVS-SM1 mediated inclusion due to their primary sequence at the exon end. Here shown as complementarity score is the number of maximal H-bonds between the U1 binding site and the U1 snRNA. (C) Violin plot showing the complementarity between the U1 snRNA and U1 binding sites of AGA-ending exons included in response to NVS-SM1 (included AGA exons) and those 5^*′*^ss matching a 5^*′*^-A_−3_ G_−2_ A_−1_ G_1_ U_2_ − 3^*′*^ motif, thus being potential candidates for NVS-SM1 mediated inclusion, but are not included (not incl. AGA Candidates). The complementarity score was calculated by summing up all H-bonds possibly formed between the two RNAs.

### Impact of U1 snRNA complementarity on compound-mediated exon inclusion

Even when exons do check all the high expression level, low baseline inclusion and exon end motif criteria, some still fail to show inclusion after compound treatment. One endogenous factor governing a 5*′* splice-site’s efficiency is the overall complementarity of this splicesite to the U1 snRNA. High complementarity enhances the recognition of the splice site, whereas low complementarity favors exon skipping (Freund et al., 2005). We therefore rationalized that complementarity to U1 snRNA may play a key role in compound-mediated splicing. Even if a modulator helps strengthen weak splice sites, a certain amount of baseline complementarity may still be required. To test this hypothesis, we calculated the maximum number of possible hydrogen bonds (H-bonds) formed between the U1 binding site of AGA exons and the U1 snRNA. Accounting for the possibility of an A_−1_ bulge formation in the pre-mRNA, we included the -4 position of the exon in the score calculation (Figure 2B). Exemplary scores for some of the exons induced by NVS-SM1 can be found in Supplementary Table S1. This score, hereafter referred to as the complementarity score, was used as a proxy for the complementarity between the 5*′*ss and U1 snRNA. For simplicity, we assumed that each potential H-bond contributes equally to the stability of the U1–5*′*ss interaction, acknowledging that individual H-bonds can differ in strength. This approach provides a tractable estimate of sequence complementarity without accounting for such variations. Analyses highlighted a significant difference between the distributions of complementarity scores for AGA exons amenable to induction by NVS-SM1 compared to non-amenable AGA exons (p-value < 0.0001, Figure 2C). We observed the same difference in analogous analyses of induced and uninduced GA exons upon Risdiplam treatment (Supplementary Figure S5A). These findings suggested that the complementarity between the 5*′*ss of target exons and U1 snRNA contributes quantitatively to exon induction upon compound treatment and is a crucial determinant of exon responsiveness to the compounds.

To verify this, we designed an experiment to modify U1 snRNA – pre-mRNA complementarity while leaving the small-molecule binding site unchanged. We ectopically expressed a U1 snRNA, harboring 6A>G and 7U>C mutations to improve pairing with the intronic region of pre-mRNAs (Figure 3A). We refer to this mutant U1 snRNA as U1-G_6_C_7_, since the changes pair with positions +6 and +7 of the exon’s U1 binding site. Whole transcriptome profiles were sequenced in biological triplicates and analyzed for exons induced by the combination of U1-G_6_C_7_ transfection and NVS-SM1 treatment at 100nM. Compared to the DMSO-treated cells transfected with U1-G_6_C_7_, 156 AGA exons were found significantly induced upon combining U1-G_6_C_7_ transfection and NVS-SM1 treatment (Supplementary Figure S1D). Of these, 29 were not detected as significantly induced upon the control combination of U1-wt transfection and NVS-SM1 treatment (FDR = 10%, dPSI >0.05).

**Figure 3.**
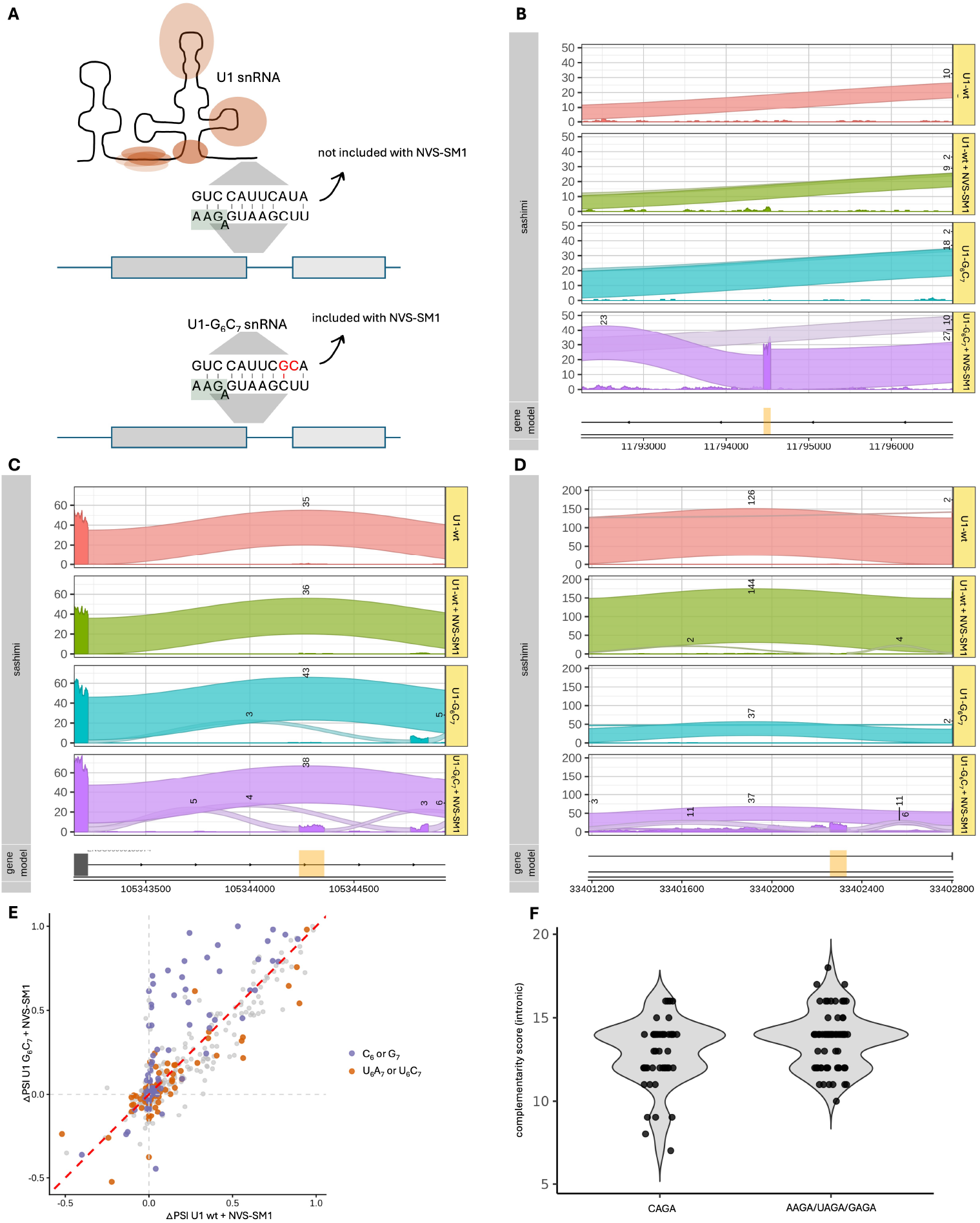
Complementarity between U1 snRNA and the U1 binding site is a critical factor for exon responsiveness. (A) Cartoon showing the complementarity of the exon highlighted in (B) with the canonical and the mutated U1 snRNA. Complementarity score with U1-wt was 19 and the score calculated for U1-G_6_ C_7_ was 22. (B) Sashimi plots of RNA-seq results deriving from HeLa cells either transfected with canonical U1 snRNA (U1-wt) or with mutated U1 snRNA (U1-G_6_ C_7_), grown in the absence or presence of NVS-SM1, each performed in triplicates. Plot highlights a cryptic exon on chromosome 3 within the gene *TAMM41*. Significant inclusion only detected in cells transfected with U1-G_6_ C_7_ and treated with 100nM NVS-SM1 (dPSI = 0.8, 10% FDR). (C) Sashimi plots of RNA-seq results deriving from the same experiment as explained in A. Plot highlights an exon on chromosome 2 within the gene *C2orf49*. Significant inclusion only detected in cells transfected with U1-G_6_ C_7_ and treated with 100nM NVS-SM1 dPSI = 0.16, 10% FDR). (D) Sashimi plots of RNA-seq results deriving from the same experiment as explained in A. Plot highlights an exon on chromosome 3 within the gene *UBP1*. Significant inclusion only detected in cells transfected with U1-G_6_ C_7_ and treated with 100nM NVS-SM1 (dPSI = 0.52, 10% FDR). (E) Scatter plot comparing exon inclusion changes (ΔPSI) upon compound treatment in cells expressing canonical (U1-wt + NVS-SM1) or mutated U1 snRNAs (U1-G_6_ C_7_ + NVS-SM1). The x-axis shows values for exons in cells transfected with canonical U1 and treated with 100 nM NVS-SM1, and the y-axis shows values for the same exons in cells transfected with U1-G_6_ C_7_ under identical treatment conditions. ΔPSI between the two conditions. Exons with higher complementarity to the mutated U1 (carrying a C at position 6 or a G at position 7 of the 5 ss) are highlighted in purple, whereas exons with higher complementarity to the canonical U1 are shown in orange. The diagonal line indicates equal ΔPSI between the two conditions. (F) Complementarity of intronic base-pairing (positions +1 through +8) of exons included upon NVS-SM1 treatment, grouped them according to their -4 position. Left group shows all exons with a C_−4_ A_−3_ G_−2_ A_−1_ sequence while the right group shows all other AGA exons (A_−4_ A_−3_ G_−2_ A_−1_, U_−4_ A_−3_ G_−2_ A_−1_ or G_−4_ A_−3_ G_−2_ A_−1_). Significant co-dependence of low-complementarity sites with the position -4 detected (p-value = 0.02).

One cryptic exon, located in the *TAMM41* gene and exclusively induced by the combination of U1-G_6_C_7_ expression and NVS-SM1 treatment, demonstrated particularly high inclusion rates, substantiated by clear exon-exon junction reads (dPSI=0.8, Figure 3B). Another cryptic exon in gene *C2orf49* was also found significantly induced upon combination treatment (dPSI = 0.16, Figure 3C). In *UBP1*, a cryptic exon demonstrated only a slight, non-significant increase with U1-wt plus NVS-SM1 (dPSI = 0.014), yet a much stronger induction with U1-G_6_C_7_ plus NVS-SM1 (dPSI = 0.52, Figure 3D). Additional examples can be found in Supplementary Figure S6A-D. Notably, exons that had a stronger complementarity with the mutant U1 snRNA, including the three examples mentioned before, demonstrated higher inclusion in response to NVS-SM1 in cells expressing the respective mutant. Whereas exons better matching U1-wt, responded stronger in cells transfected with the canonical U1 snRNA. These trends were reflected in both when comparing ΔPSIs (Figure 3E) and absolute PSIs between cells transfected with the different U1 snRNAs in combination to the compound treatments (Supplementary Figure S7A).

To test whether this observation applies to other splicing modifiers as well, we repeated the experiment using 1µM of Risdiplam. We compared U1-G_6_C_7_–transfected cells treated with 1µM of Risdiplam to transfected cells treated with DMSO as a control and found 106 significantly induced GA exons (Supplementary Figure S1E). Of these, 23 exons were not significantly induced upon the combination of U1-wt transfection and 1µM Risdiplam. As previously observed for NVS-SM1, this set of exons demonstrated a shift to higher complementarity scores with U1-G_6_C_7_ compared to canonical U1 snRNA (Supplementary Figure S5B-F and S7B). Overall, these data show that baseline complementarity between U1 snRNA and the target exons contributes to compound-mediated exon induction.

### Role of the -4 exon end position in compound-mediated splicing

In an attempt to unify the A_−1_ bulge formation model (Palacino et al., 2015; Campagne et al., 2019) and our observations on complementarity requirements, we tested how the -4 nucleotide at the exon’s 3*′* end affects NVS-SM1 and Risdiplam activity. As illustrated in Figure 2B, a cytosine at position -4 could pair with a guanine in position 11 of the U1 snRNA, assuming an A-1 bulge forms in the pre-mRNA, and thus restoring the canonical 5*′*ss initiation sequence CAG. To test this model in a cellular context, we analyzed the 125 AGA exons induced by NVS-SM1 in SH-SY5Y cells and grouped them according to their -4 position, resulting in 50 exons with a 5*′*-CAGA-3*′* ending and 75 exons with either UAGA, GAGA or AAGA ends. Strikingly, when focusing on the complementarity of the intronic base-pairing (positions +1 through +8) to a canonical U1 snRNA, we found that low-complementarity sites favor a cytosine at position -4 (p-value = 0.02, Figure 3F). This suggests that base-pairing at position -4 can help compensate for weak U1 binding in the intron.

To experimentally test the influence of the -4 position on the formation of an RNA duplex with a A_−1_ bulge, we employed a Fluorescence Polarization (FP) assay based on a TAMRA-labeled 16 nucleotide RNA representing the U1 snRNA sequence. With increasing concentrations of *HTT* pseudoexon 5*′*ss, we observed an increase in fluorescence polarization, indicating binding of the labelled U1 sequence (EC_50_ = 252nM, Figure 4A/B). Repeating the experiment with a target RNA harboring a C>G mutation at the -4 position (-4C>G, Figure 4B), we observed a significant decrease in FP signal compared to the wt RNA (EC_50_ = 627nM, Figure 4A). This finding corroborates the influence of the nucleotide at position -4 on the overall base pairing between U1 snRNA and 5*′*ss, which can compensate for weak intronic U1 snRNA basepairing.

**Figure 4.**
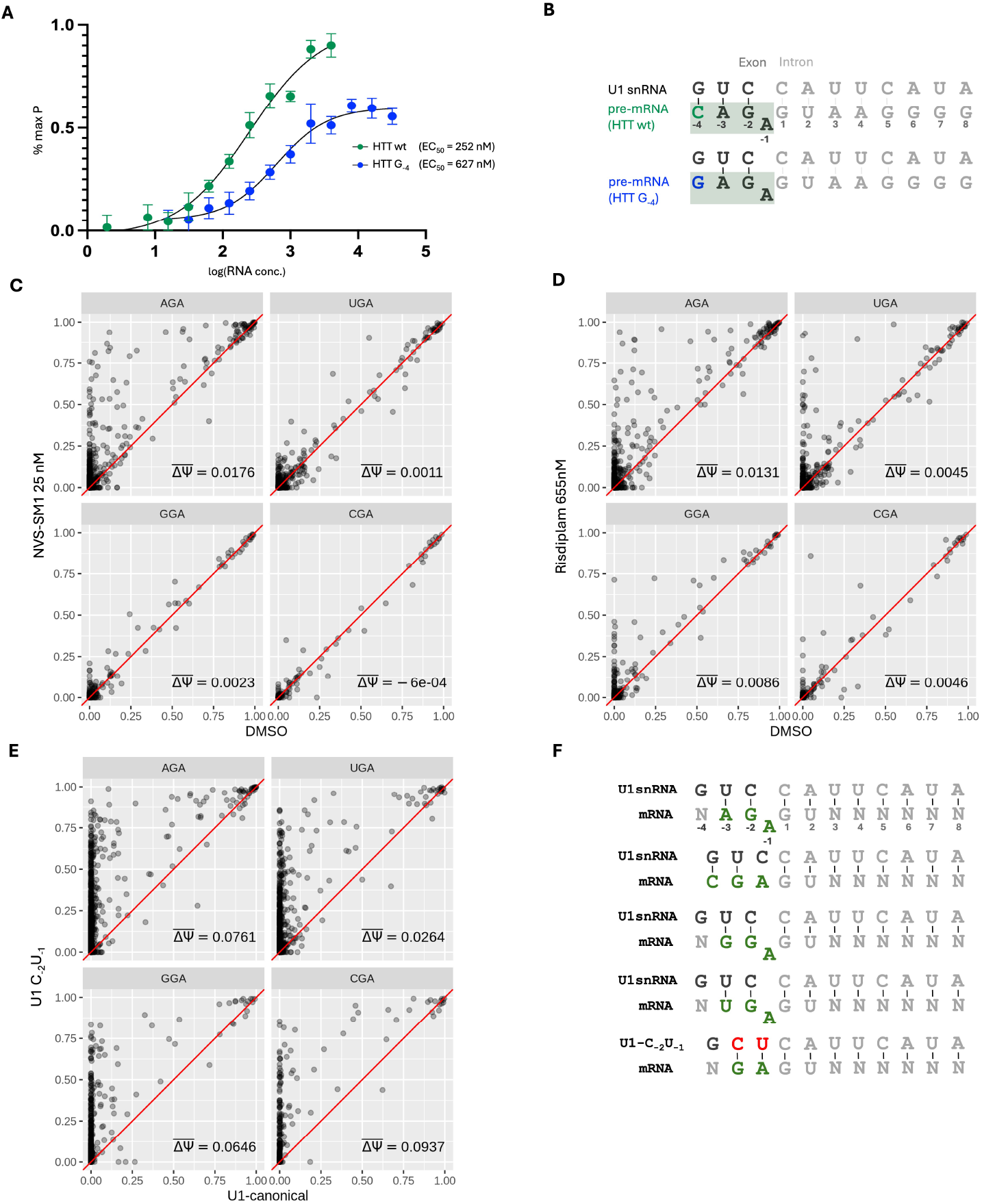
Bulge formation and sequence features of exons induced by NVS-SM1 and Risdiplam. (A) Fluorescence polarization (FP) binding curves of TAMRA-labeled U1 snRNA incubated with serial dilutions of the U1 binding site of the *HTT* pseudoexon, either the wild-type sequence (wt), or a mutated version (HTT G_−4_). Samples were measured in triplicates, and the data analyzed in GraphPad Prism 9.0. A trendline was generated using a sigmoidal dose-response curve fit to determine an EC_50_ value. (B) Constructs used for the FP-assay. (C-E) Scatter plots showing PSI values of GA-ending exons, grouped according to their –3 position (AGA, UGA, GGA, or CGA). The red diagonal line indicates equal PSI values between the compared conditions and ΔΨ the mean change in inclusion. (C) PSI values in cells treated with 25 nM NVS-SM1 or DMSO. (D) PSI values in cells treated with 655 nM Risdiplam or DMSO. (E) PSI values in cells transfected with either U1-C_−2_ U_−1_ or canonical U1 snRNA. (F) Schematic of potential bulge formation in RNA duplexes between U1 snRNA and the exon–intron junction of GA-ending exons. U1-C_−2_ U_−1_ shows sequence of the mutated U1 snRNA, modified to directly induce GA-ending exons in the absence of a compound.

### Primary sequence features of GA exons induced by NVS-SM1 and Risdiplam

Considering the influence of the -4 position on the pairing between U1 snRNA and non-canonical GA exons, we analyzed the primary sequence features of exons induced by NVS-SM1. In line with the established GAspecificities of these compounds, we observed an enrichment for GA exon ends (positions -2 and -1) in the 5*′*ss motifs of NVS-SM1 and Risdiplam responsive exons in SH-SY5Y cells. In NVS-SM1 treated cells, the A_−3_G_−2_A_−1_ motif was enriched (Figure 1C). Analyzing the nucleotide distribution at position -2 of all induced A-ending exons, we found a near-exclusive occurrence of guanine (91%, i.e. 146 out of 160 upon NVS-SM1 treatment and 94%, i.e. 151 out of 161 upon Risdiplam treatment) (Supplementary Figure S4A). Consistent with the observed motif, a robust enrichment for adenine in response to NVS-SM1 could be detected at the -3 position of induced GA-ending exons (86%, i.e. 125 out of 146). This motif preference correlated with exon responsiveness upon compound treatment, reflected by overall PSI values. AGA-ending exons are most efficiently induced by NVS-SM1, while CGA-ending exons, intriguingly, did not show any average inclusion changes(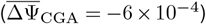) (Figure 4C). This pattern is consistent with the differential exon inclusion analysis, which revealed that NVS-SM1 induces several GGA and UGA exons (14 and 7, respectively) but no CGA exons (Supplementary Figure S4B).

To explore whether CGA exons can in principle beinduced, we transfected HeLa cells with a modified U1 snRNA harboring -2U>C and -1C>U mutations, designed to stabilize all GA-ending exons (U1C_−2_U_−1_; Figure 4F). Transcriptome sequencing of transfected cells identified exons specifically induced by U1-C_−2_U_−1_ compared to those induced by canonical U1 snRNA (Supplementary Figure S1F). Under these conditions, CGA exons show increased inclusion comparable to the effect on other GA-ending exons, with the highest mean change among them (Figure 4E,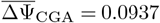). The differential analysis revealed inclusion of 96 CGA exons (9% of all induced GA-ending exons; Supplementary Figure S4E). In contrast, when transfecting cells with the canonical U1 snRNA and treating cells with NVS-SM1, inclusion of CGA-ending exons is almost absent, with only one detected exon (less than 1% of induced GA-ending exons; Supplementary Figure S4E). Similar to Branaplam, also Risdiplam disfavored CGA-ending exons (Figure 4D and Supplementary Figure S4B). These findings confirm that CGA exons are generally amenable to splicing modulation when U1 binding is stabilized but are refractory to small-molecule–mediated activation under native conditions. A likely explanation is that the C_−3_G_−2_A_−1_ sequence prevents formation of an A bulge due to the energetically favorable G–C pairing at position -3 (Figure 4F). This, in turn, suggests the bulge stabilization at position -1 to not just be the result of compound binding, but actually a prerequisite for it. To further test this model, we transfected cells with an U1 snRNA variant (U1-C_−2_G_−1_) that disfavors bulge formation with AGA exons. U1-C_−2_G_−1_ expression moderately reduced the splicing response to NVS-SM1, further supporting that compound activity in the spliceosome depends on the ability to form an 1-A bulge (AGA exons with dPSI>0.05 in U1wt, U1-C_−2_G_−1_ showed a mean dPSI reduction of 0.040) (Supplementary Figure S4F).

As mentioned, with NVS-SM1 treatment, AGA exons showed the strongest increase in inclusion compared to DMSO controls. Intriguinly, for GA exons lacking an adenine at position –3, we noticed a global compound response only when an adenine was present at position –4 (A(C|G|U)GA) (Figure 5A). In contrast, GA-ending exons without adenine at either positions –4 or–3 ((C|G|U)(C|G|U)GA) display almost no change in inclusion(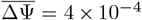, Figure 5A). Analyses of sequence composition of statistically significantly included exons supported these observations: while induced AGA exons exhibit an even nucleotide distribution at position –4 (Supplementary Figure S4C), significantly induced GGA and UGA exons almost exclusively feature an adenine at position -4 (Supplementary Figure S4D). Risdiplam showed the same pattern, where only GA-ending exons containing a second adenine at either position -3 or -4 also showed a response to the compound (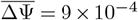, Figure 5B and Supplementary Figure S4C). The strong preference for AGA-, AGGA-, and AUGA-ending exons thus reflects a broader requirement for a secondary adenine at positions –3 or –4. As adenines in position –4 are unlikely to contribute to U1–5*′*ss duplex stabilization (Figure 4F), their presence may favor compound binding.

**Figure 5.**
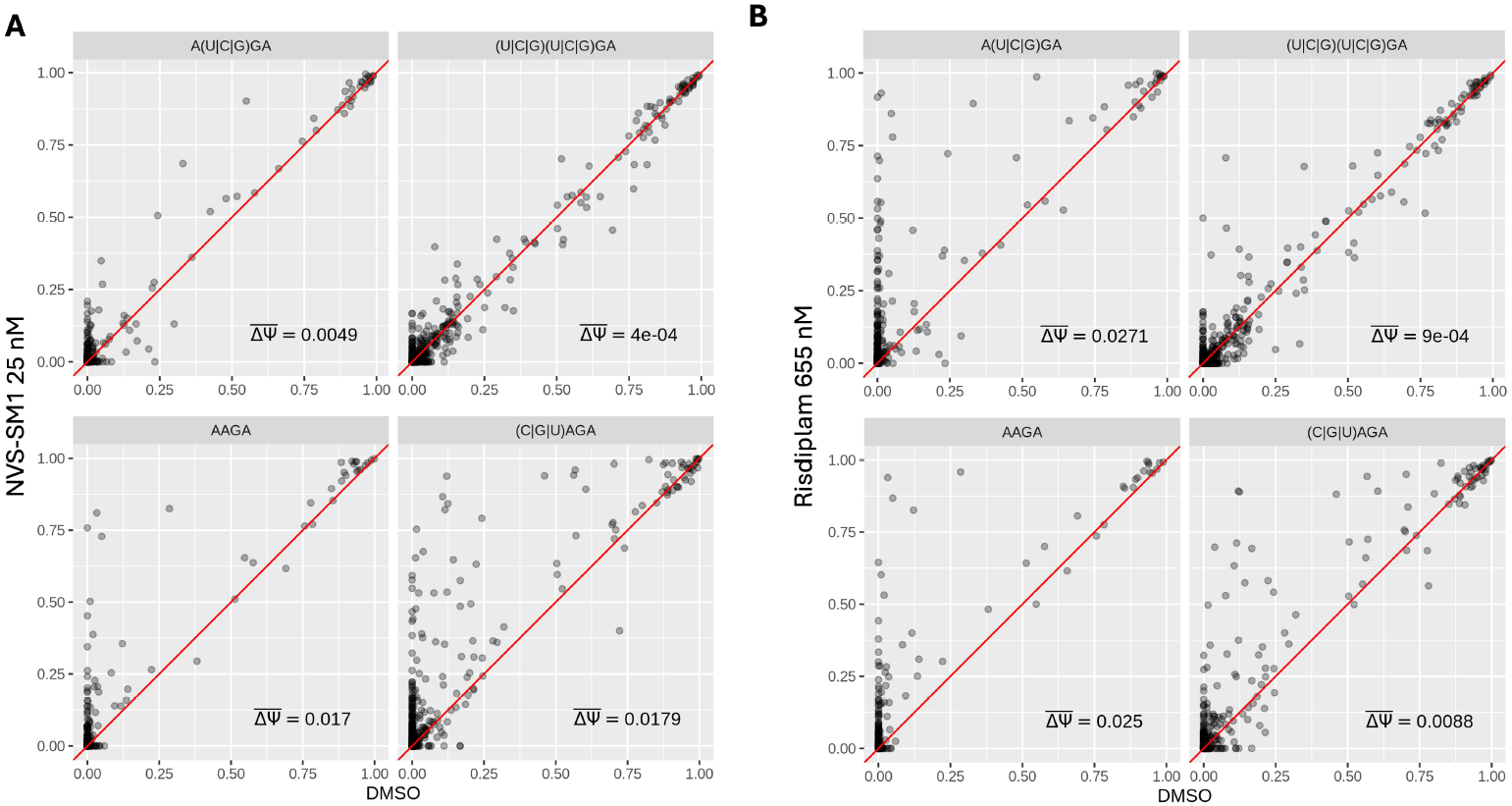
Necessity of an A at -3 or -4 position. Scatter plots showing PSI values of GA-ending exons in cells treated with a compound or DMSO. Exons are grouped based on the presence of adenine at the –3 and –4 positions: A at both positions (AAGA), A only at –3 ((C|G|U)AGA), A only at –4 (A(C|G|U)GA), or A absent at both positions ((C|G|U)(C|G|U)GA). The red diagonal line indicates equal PSI values between the compared conditions. The mean PSI change ΔΨ is shown for each panel. (A) Cells treated with 25 nM NVS-SM1. (B) Cells treated with 655 nM Risdiplam.

### NVS-SM1 interaction with A-1 bulge in RNA duplexes

To understand the dynamics of A_−1_ bulge structure stabilization, we designed a 13 nucleotide RNA probe containing the 5*′*ss sequence of the *HTT* pseudoexon (positions -4 to +9), in which we replaced the adenine at position -1 with the fluorescent analogue 2-aminopurine (2-AP). The fluorescence of 2-AP depends on its microenvironment and, while un-impacted by events such as hydrogen-bond formation, is being quenched by base-stacking and clashes with neighboring nucleobases (Rachofsky et al., 2001). Therefore, 2-AP can act as a sensitive reporter to monitor changes to the secondary or tertiary structure of RNA, when incorporated into a nucleotide strand (Rau and Hall, 2015). Moreover, those characteristics can be applied to detect and monitor ligand-engagement to RNA (Bradrick and Marino, 2004; Chen et al., 2020). Thus, the 2-AP probe enabled monitoring both of A_−1_ bulge secondary structure formations. We incubated the 2-AP probe with wildtype U1 snRNA or modified U1 snRNA oligonucleotides, respectively. The modified U1 oligo, harboring the mutations -3G>U, -2U>C and -1C>U, reverse complements the 2-AP probe and thus forms an RNA duplex without any mis-match induced secondary structures. Hence, we termed this modified U1 snRNA oligo “closed U1” (Figure 6A).

**Figure 6.**
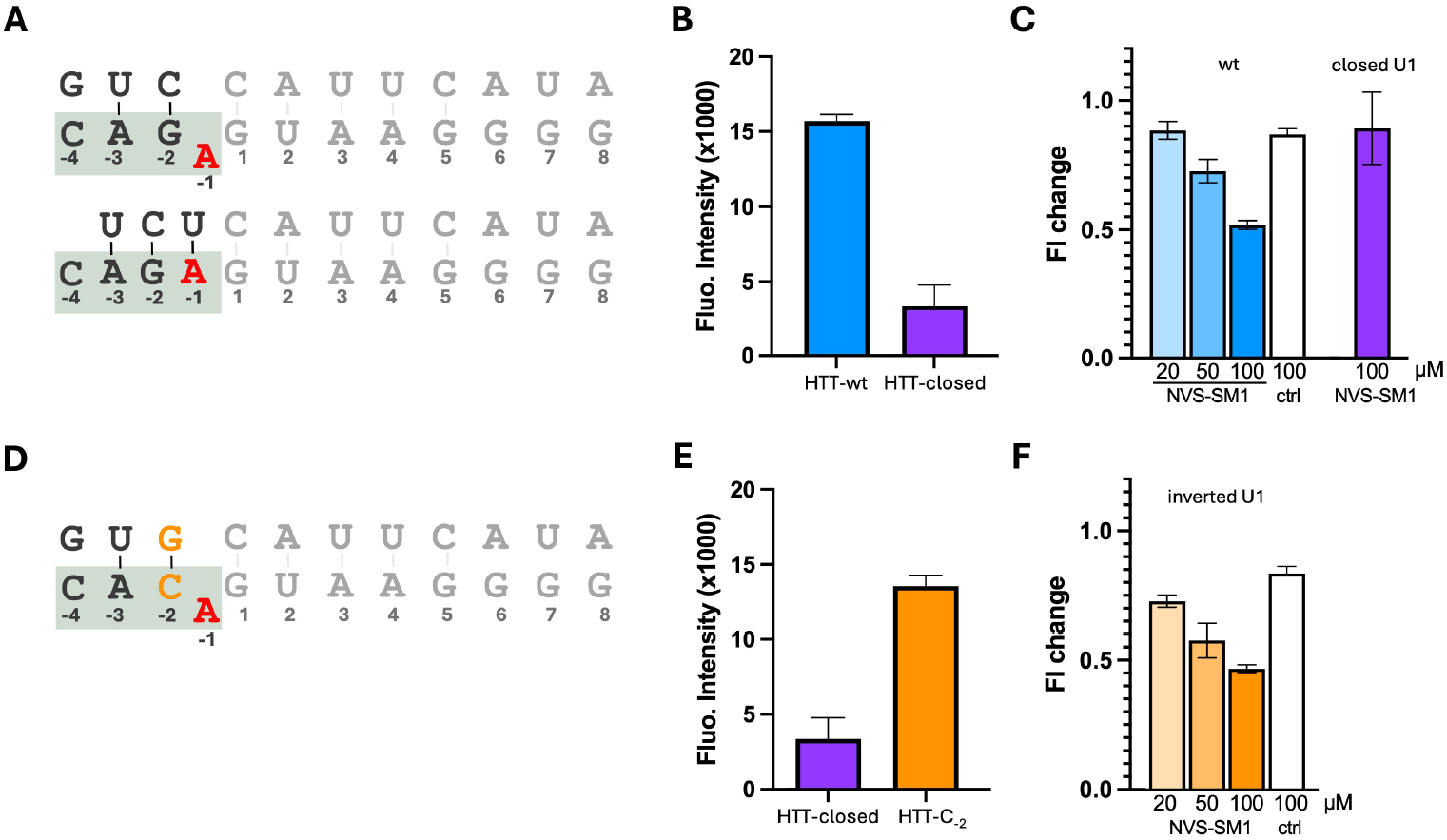
Biochemical Assays to investigate bulge formation at U1 binding site of the *HTT* pseudoexon. (A) RNA constructs used for the 2-AP assay shown in (B) and (C). RNA constructs carrying either wild-type sequence of the RNA complex formed when *HTT* pseudoexon is recognized (wt, top) or a mutated version, forming a closed duplex (bottom, closed U1). Position of the 2-AP mutation highlighted in red. (B) Fluorescence Intensity (FI) measured in the 2-Aminopurine (2-AP) assay with wt or closed U1 RNA construct (sequences shown in (A)). (C) Changes in FI upon titration of NVS-SM1 or inactive compound (ctrl, shown as neg.ctrl in Supplementary Figure S1G) to RNA duplexes, relative to the DMSO condition. (D) RNA constructs used for the 2-AP assay shown in (E) and (F). RNA construct carries a mutated version of the RNA complex formed when *HTT* pseudoexon is recognized, changing primary sequence while maintaining possibility of bulge formation. 2-AP mutation highlighted in red. (E) Fluorescence Intensity (FI) measured in the 2-Aminopurine (2-AP) assay with the closed U1 or inverted U1 RNA construct (sequences shown in (A) and (D)). (F) Changes in FI upon titration of NVS-SM1 or inactive compound (ctrl, shown as neg.ctrl in Supplementary Figure S1G) to RNA duplexes, relative to the DMSO condition.

As expected, adding the closed U1 construct caused strong quenching of 2-AP fluorescence, indicating base stacking in the RNA duplex (Figure 6B). Conversely, when adding the canonical U1 RNA oligo, the 2-AP fluorescence was significantly higher compared to the closed construct, indicating that the 2-AP base was not stacked in the RNA duplex, thus confirming the presence of a bulge structure (Figure 6A). Next, we monitored the 2-AP fluorescence intensity (FI) changes in the presence of canonical U1 RNA oligo, with increasing concentrations of either NVS-SM1 or a structurally related, splicing-inactive compound (ctrl; Supplementary Figure S1G). While the inactive compound did not cause any fluorescence decrease, NVS-SM1 caused a concentration-dependent quenching of the 2-AP FI, confirming its interaction with the A_−1_ bulge in the RNA duplex (Figure 6C).

### Primary and secondary RNA structures define the splicing modulator target exon space

To better understand how RNA structure and sequence affect compound binding, we adapted the 2-AP assay to test whether NVS-SM1 could bind the same secondary structures in the context of different primary sequences. Therefore, we modified the HTT 5*′*ss 2-AP probe with a - 2G>C mutation to mimic an ACA-ending exon. To match this oligo probe, we designed a 12 base oligonucleotide with the U1 snRNA sequence harboring a -2C>G mutation, called “inverted U1” (Figure 6D). The combination of both the modified HTT 5*′*ss probe and the inverted U1 oligo allows for and favors the formation of a bulged-out 2-AP reporter nucleotide (Figure 6C/F). The baseline fluorescence intensity of the HTT-C_−2_ RNA complex was found similar to the FI observed for the HTT-wt complex (Figure 6F), suggesting a resembling formation of a bulged-out adenine reporter nucleotide at position - 1 of the RNA duplex. Subjecting this reporter complex to increasing concentrations of NVS-SM1, we observed a dose-dependent quenching of the 2-AP probe, this time embedded into a different sequence context (Figure 6F). These data demonstrate that binding of NVS-SM1 to adenine bulge structures occurs independent of the primary RNA sequence. This observation, in line with our genetic perturbation experiments, highlights the effect of the highly conserved U1 snRNA sequence as a major determinant, governing the specificity of small molecule splicing modulators, such as NVS-SM1.

### Contribution of regulatory elements

We further characterized the contribution of regulatory elements, beyond the U1 snRNA binding sites, to compound activity. Analyses of the abundance of predicted exonic splicing enhancer (ESE) and exonic splicing silencer (ESS) motifs showed no significant differences between induced AGA exons and non-induced AGA exons. To integrate broader regulatory information, we trained a gradient-boosted decision tree model (XGBoost, Figure 7A, B) to classify AGA exons between inducible and non-inducible using sequence context and motif predictions. Overall, Shapley additive explanations values (SHAP) of the model confirmed our previous observations on the contribution of the different U1 snRNA positions to compound mediated exon induction (Figure 7C).

**Figure 7.**
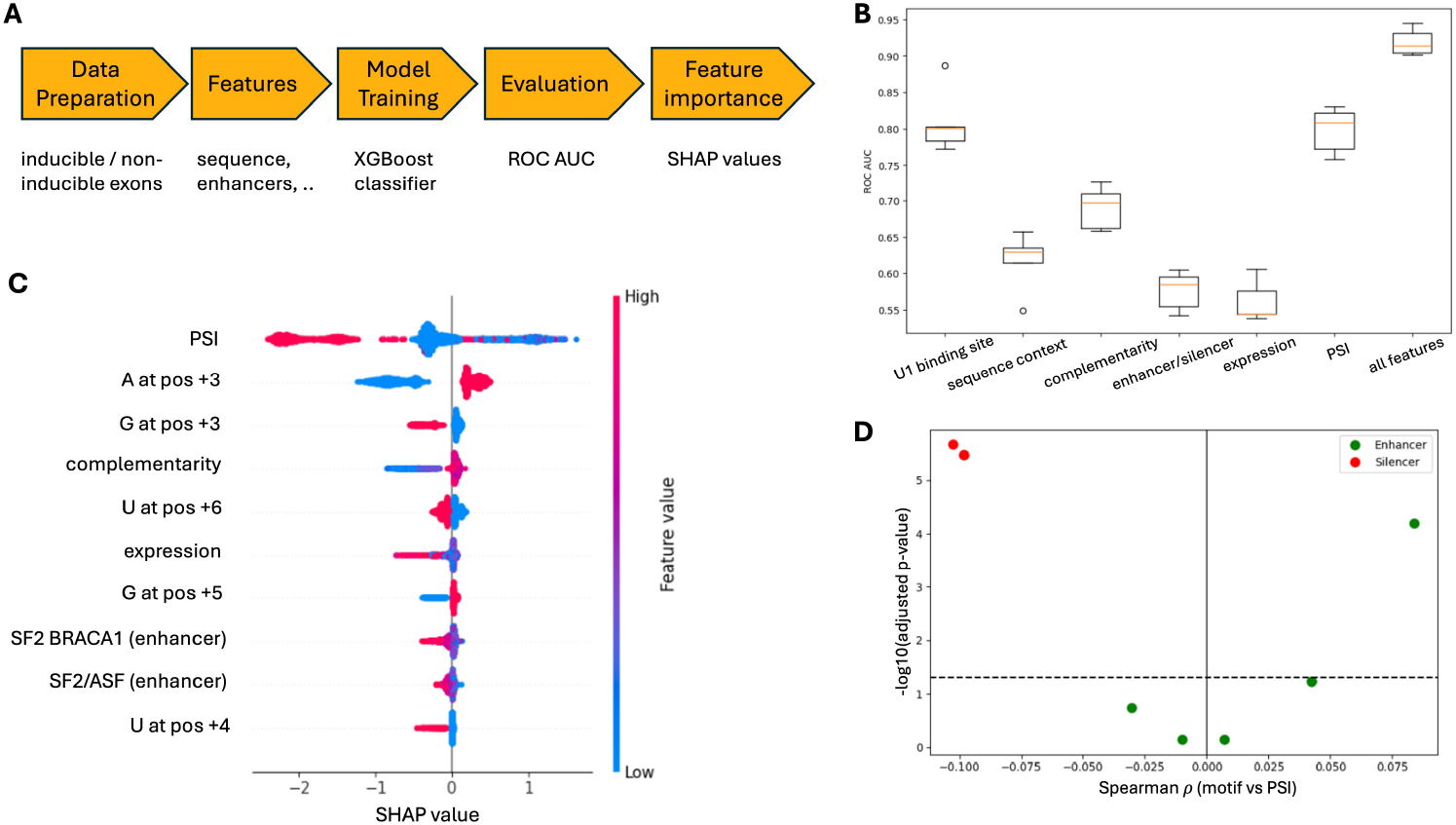
Machine-learning model and contriutions of different features on prediction. (A) Schematic overview of the machine-learning workflow. The model integrates sequence features, enhancer/silencer scores and expression information to distinguish inducible AGA exons from non-inducible AGA exons. An XGBoost classifier was trained and evaluated using cross-validated ROC AUC, followed by SHAP-based feature interpretability. (B) Comparison of model performance across different feature sets using 5-fold cross-validated ROC AUC. Each boxplot represents the distribution of AUC values across folds for one feature set. (C) SHAP summary plot displaying the top features contributing to the model predictions. Each point represents one splice site. The horizontal location shows whether the effect of that value is associated with a higher or lower prediction and the color represents the original value of the feature. (D) Volcano plot showing the correlation test between silencer and enhancer motif abundance and basal PSI in DMSO samples. Each point represents an enhancer or silencer motif, colored by category. The x-axis displays the spearman correlation coefficient (ρ), and the y-axis shows the –log10 adjusted p-value. Horizontal and vertical reference lines indicate the significance threshold (padj = 0.05) and zero correlation, respectively.

Moreover, a model based solely on enhancer and silencer predictions performed only marginally above random (Figure 7B), confirming that these predictions alone provide a limited explanatory power. In contrast, a model trained on the surrounding sequence context, while explicitly excluding the U1 snRNA binding site, achieved a ROC AUC above 0.6 (Figure 7B), demonstrating that additional regulatory information embedded in the flanking sequence contributes to compound responsiveness. Notably, a model restricted to the U1 binding site itself (positions -4 to +8) performed even better, reaching a ROC AUC of 0.8 (Figure 7B). This underscores that this region remains the dominant determinant of responsiveness, while the broader sequence context provides secondary but meaningful regulatory cues. The SHAP analysis identified baseline inclusion (PSI in DMSO) as one of the most influential features. Specifically, AGA exons with a high baseline PSIs were not further induced by compound treatment. Interestingly, we found that baseline PSI was significantly correlated with the number of ESE and anticorrelated with the number of ESS (Figure 7D).

Overall, the interpretation of this model aligns with our previous analyses presented in this study, which highlighted the several key characteristics, such as complementarity with the U1 snRNA. Further, these results suggest that compound responsiveness is also shaped by subtle sequence embedded features and integrated regulatory effects.

## Conclusion

This study provides comprehensive insights into the mechanisms underlying the specificity of small molecule splicing modifiers, particularly focusing on NVS-SM1 (Branaplam) and its interaction with the splicing machinery. Although these compounds target exons with specific exon-end, not all of these exons are equally responsive. This suggests that additional determinants governing compound specificity are required to render an exon responsive to small molecule splicing modifiers. By integrating transcriptomic profiling, biochemical assays, U1 perturbation experiments, regulatory motif annotation, and machine-learning-based feature interpretation, we identify multiple layers of regulation that together determine whether an exon can be modulated by small molecules.

First, we highlighted that the landscape of inducible exons is shaped significantly by cell type specific expression, alongside with primary sequence. Furthermore, we analyzed the contributions of the downstream intronic nucleotides within the U1 snRNA binding site of the target exon. Their functional importance has remained largely unexplored in the context of smallmolecule splicing modulation, particularly across different exon architectures. It has not only remained unclear whether these positions consistently influence compound responsiveness across different exons, but also how they contribute mechanistically to modulator activity. Our investigation into their complementarity between the 5*′*ss and the U1 snRNA revealed a crucial role that the U1 snRNA plays in mediating compound-mediated exon inclusion. Exons with higher complementarity proved more likely to be targetable by splicing modifiers, whereas those with lower complementarity tend to be skipped, regardless of the intervention. Our experiments with mutated U1 snRNA showed that changing U1–5*′*ss pairing can reprogram compound specificity. This, in turn, highlights the strong dependence of the observed compound specificity on the highly conserved U1 snRNA sequence and explains why the actual specificity of these compounds tends to exceed theoretical expectations.

To gain a deeper understanding of what governs the apparent primary sequence specificity of NVS-SM1 and Risdiplam, we took a closer look at potential bulge formation at the -1 position of the exon end. In line with previously reported biophysical data (Malard et al., 2024), our cellular and biochemical data corroborates the necessity of a bulge formation at the -1 position of the exon end. In other biological contexts, covariation helps to identify RNA structural specificity; however, in this case, covariation does not occur due to the high conservation of the U1 snRNA sequence. Thus, we used mutated U1 snRNA in our assays to test if we could shift the sequence specificity of the compounds. Notably, we were able to induce compound binding to a previously non-amenable CA-ending exon when the U1 snRNA was mutated to maintain a bulged-out adenine at the -1 position of the pre-mRNA 5*′*ss. Also, our transcriptomic analyses showed that exons lacking the ability to form a bulge at the -1 position (CGA-ending exons), even if they fulfill the N_−3_G_−2_A_−1_ motif, are refractory to compound modulation. Furthermore, expressing a U1 snRNA variant that disfavors this bulge formation with AGA exons reduces the effect of NVSSM1 on AGA exons. These findings further emphasizes that the formation of the required U1-5*′*ss RNA duplex secondary structure is not a merely the result of compound binding, but rather a prerequiste for compound binding.

Consistent with observations by Ishigami et al. (Ishigami et al., 2024), we also identified that the –4 position as a critical determinant of compound responsiveness. Our results corroborates that this position substantially influences drug activity and provide an explanation for why many N_−3_G_−2_A_−1_-ending exons fail to be induced. For AGA exons, the presence of a C in the -4 position compensates for low U1 snRNA intronic complementarity. Furthermore, our findings lead us to propose that the AGA or ANGA sequence specificity arises from two key factors. Firstly, it is essential to maintain a specific secondary structure that allows for bulge formation, resulting in the requirement of a G at the -2 position. Secondly, two adenines appear to be essential, one at position -1 and one at either position -3 or -4. Since an adenine at position –4 is unlikely to contribute to U1–5*′*ss duplex stabilization, these adenines are likely forming interactions with the compounds, thereby contributing to the observed primary sequence specificity. Together, these results define primary sequence codependencies that shape bulge formation and the surrounding U1–5*′*ss duplex architecture as the primary determinants of NVS-SM1 and risdiplam responsiveness.

However, the presence of unresponsive exons that do match our identified characteristics of inducibility suggests additional features and regulatory elements. Previous publications have highlighted the importance of such elements. For example, the purine-rich element in SMN2 exon7 (Tang et al., 2021), have been shown to shape compound binding. Further work on extended exonic sequence similarly shows that local context can influence drug activity (Campagne et al., 2024). Although our genome-wide analyses did not reveal any motif beyond the U1 binding site or splicing enhancers that were broadly present in compound-induced exons, we can not exclude that individual regulatory elements may still modulate responsiveness in specific exons. Indeed, in our machine learning model, the feature importance analysis highlighted a contribution of local sequence context to compound modulation. However, baseline AGA exon inclusion in the absense of a compound was correlated with the number of ESE and anticorrelated with the number of ESS, suggesting that these regulatory elements may compensate for AGA weak splice sites under normal physiological conditions. Overall, our findings shed light on the intricate interplay between primary sequence and secondary structure requirements in governing the overall observed specificity of small molecule splicing modifiers i n c ells. Understanding this is essential to decipher the rules governing compound specificity, improve predictiv models for target exons, and potentially tune the activity of splicing modulators in a controlled manner. On one hand, the specificity o f a U 1 s nRNA c an b e i nfluenced by the addition of a splicing-modulating compound. Genetic manipulation of the U1 snRNA, on the other hand, can change the specificity of a splicing modulating compound. These insights may aid future development of small molecules targeting RNA splicing as well as RNA secondary structures. Moreover, these findings will help to refine prediction algorithms, paving the way for identification of novel targets and rational design of new splicing-modifying compounds.

## Methods

**Cell Culture:** SH-SY5Y neuroblastoma cells were cultured using Advanced MEM media supplemented with 10% heat-inactivated fetal bovine serum (HI FBS) and 1X Glutamine (5 mL per 500 mL of media). Upon reaching confluency, the cells were sub-cultured and plated into 12-well plates at a density of 0.6 x 106 cells/mL, with 1 mL per well. On the second day post-plating, the cells were treated with compounds at different concentrations, DMSO treatment condition is used as control. Cells were harvested 24 hours posttreatment. For iNGN2 cells, day 3 iNGN2 neurons were thawed and plated onto six-well plates, pre-coated with poly-D-lysine (PDL) and a mixture of 1X Matrigel and laminin (1:300, Sigma-Aldrich, L2020-1MG). The coating procedure involved incubating the plates at 37°C for at least 1 to 1.5 hours prior to cell plating. Neurons were maintained by providing half of the usual volume of culture medium every other day until day 10. On day 10, they were treated with compounds at different concentrations. DMSO treatment condition is used as control. Cells were harvested 24 hours posttreatment. HeLa 293 cells were maintained in DMEM with 10% fetal bovine serum (FBS) and 1% GlutaMax at 37 °C with 5% CO_2_. They were plated on a 24-well plate (80.000 cells/well) and treated with potential compounds or DMSO as control the next day. Cells were harvested 24 hours post-treatment.**Expression of U1 snRNA:** For the ectopical expression of U1 snRNA, we used a plasmid either carrying the sequence of the canonical U1 snRNA or a mutated U1 snRNA version (e.g. U1-G_6_C_7_). HeLa cells were plated on a 24-well plate (80.000 cells/well) and transfected the following day with 0.3 µg DNA and 0.9 ml LipofectamineTM 2000 (Thermo Fisher Scientific), according to the supplier’s protocol. After 24 h, medium was changed and potential compounds added or DMSO as a control. After 24 h incubation, RNA purification was performed using the *Quick* -RNA Miniprep Kit (Zymo Research).**NMD inhibition:** We inhibited major RNA decay pathways in SH-SY5Y cells by depleting XRN1 and EXOSC4. The culture medium was aspirated and replaced with siRNA delivery medium containing Accell siRNA targeting XRN1 and EXOSC4 or with a non-targeting control (NTC). Cells were incubated for 72 hours to achieve gene knockdown. Following this incubation, either DMSO or the splicing compound was added to the cells for an additional 24 hours. Knockdown efficiency was verified by qRT-PCR. **RNA sample preparation and sequencing:** RNA samples were collected from SH-SY5Y and iNGN2 cells treated with DMSO or compounds using the RNeasy Plus 96 Kit (Qiagen) according to the manufacturer’s instructions. RNA purification from HeLa cells purification was performed using the *Quick* -RNA Miniprep Kit (Zymo Research). All RNA-seq experiments were performed in biological triplicates. RNA quality was assessed using the TapeStation 4200, with all samples exhibiting an RNA integrity number (RIN) greater than 9. Next-generation sequencing libraries were prepared with the TruSeq Stranded mRNA Sample Preparation Kit (Illumina) from 200 ng of input RNA, following the manufacturer’s protocol. The libraries were then pooled and loaded onto a NovaSeq 6000 (Illumina) for paired-end 51 bp sequencing, yielding an average of 30-40 million reads per sample. Quality control analyses confirmed high reproducibility among biological replicates and clear separation between treatment groups. Principal component analysis (PCA) of exon inclusion levels demonstrated clustering by treatment condition (Supplementary Figure S8 and S9), and pairwise sample–sample correlations exceeded 0.95 across replicates (Supplementary Figure S10, S11, S12, S13, S14 and S15), confirming data consistency prior to downstream analyses. **Detection of novel exons and compound-mediated splicing:** RNA-seq data were mapped to the hg38 human reference genome using STAR. To identify unannotated exons in the RNA-seq data, we used the split alignments reported by STAR. For a given RNA-seq sample, we mapped introns based on reads whose mapping to the reference genome was split between non-consecutive nucleotides (split reads). For two consecutive annotated exons, these reads would partially map to one exon and partially to the second exon, thus leaving an unmapped region in between that corresponds to the intronic coor-dinate. As the mapping of all intronic regions of a gene defines the coordinates of the non-terminal exons of a gene, these regions were considered potential novel exons. We considered introns for a given sample when they were supported by at least 2 reads and filtered for novel exons that were supported by at least one third of the samples in each experiment. These potential novel exons, if unannotated, were included to the reference transcript annotation for downstream analyses. For each exon in each sample, we counted the number of reads whose mapping supported the inclusion of the exon and the number of reads whose mapping supported the exclusion of the exon. We estimated percent spliced in values as described in (Schafer et al., 2015) and used the Bioconductor package DEXSeq to test for differences between conditions as described in (Reyes and Huber, 2018). **FP binding assay:** In the FP binding experiments, 20 nmol/L TAMRA-labeled U1 snRNA target oligonucleotides were incubated with increasing concentrations of probe oligonucleotides carrying the sequence of the U1 binding site of the pseudoexon in the *HTT* gene that is being included in response to NVS-SM1. 50 mM Tris-HCl (pH 7.4), 5 mM MgCl_2_, 75 mM KCl and 0.05% Tween were used as buffer. The samples were thoroughly mixed, incubated for 5 min at 95 °C to unfold possible RNA structures and slowly cooled down to room temperature (RT). Splicing modifiers were added and incubated for another 1 h at RT. In white 384-well microplates with 30 µL per well using a PHERAstar, the parallel emission intensity (S) and perpendicular emission intensity (P) were measured. FP values were calculated using the following equation, as described previously (Zhang et al., 2006; Zhu et al., 2018): *FP* (*mP*) = 1000 ∗ (*S* − *G* ∗ *P*)*/*(*S* +*G* ∗ *P*) All samples were measured in triplicates. The FP values were plotted against the concentration of the probe RNA oligonucleotide in each experiment. Using GraphPad Prism 9.0, a sigmoidal dose-response curve was added to determine EC_50_ values. **2-AP assay:** Oligonucleotides carrying the sequence of the exon-binding part within the U1 snRNA and those with sequences corresponding to the U1 binding site that contain a 2-AP substitution at the -1 position, were mixed in a 1× Assay Buffer (0.185 M NaCl, 8 mM NaH_2_PO_4_, pH 6.3). In the absence or presence of potentially binding compounds, they were heated to 70 °C for 3 min and slowly cooled down to allow folding. In the absence of a compound DMSO is used as control. Fluorescence excitation and emission wavelengths (310nn and 390nm, respectively) were measured in a 384-well plate at RT with a Tecan plate reader (gain, 155; integration time, 20 µs). All assays were carried out in triplicates. **Analysis of exonic splicing enhancers and silencers:** ESEs were identified using PSSMs for the SR-protein motifs SF2/ASF, SF2/ASF (IgM/BRCA1 variant), SC35, SRp40, and SRp55 (Liu et al. (1998); Cartegni et al. (2003)). For each exon the last 50nt were scanned, and scored by summing nucleotide-specific PSSM weights. For each motif, we recorded the number of scoring above the motif’s canonical threshold, as well as the maximum and mean PSSM score across the exon. ESS content was assessed by scanning for hexamer motifs belonging to the published FAS-hex2 (176 hexamers) and FAS-hex3 (103 high-confidence hexamers) datasets (Wang et al. (2004)). For each exon, we computed the density ESS hits within the last 50nt. **Machine-learning classification of induced exons:** To predict exon inducibility, we trained gradient-boosted decision tree models (XGBoost) using different feature groups. For each feature set, we performed 5-fold stratified cross-validation and computed ROC AUC values. Feature vectors were generated by flattening all list- or tensor-based entries into numerical arrays. Hyperparameter optimisation identified a best-performing model with regularisation, subsampling, and class-imbalance correction.

## Data Availability

The RNA-seq data generated in this study have been deposited in the NCBI Sequence Read Archive (SRA) and will be available upon journal publication.

## Code Availability

All scripts to reproduce the analyses and figures will be available in a GitHub repository upon journal publication.

## Acknowledgment

We thank everyone involved in projects working with splicing modifiers. Moreover, we want to thank Gregory Marszalek for his support in the lab and Anne Granger for her support through the postdoctoral program at Novartis.

## Author Contributions

F.L. designed and performed most experiments; carried out analyses; and wrote the manuscript. J.K. performed library preparation and sequencing. M.A. supported sequencing experiments. A.B. contributed to analysis of sequencing experiments and exon selection. W.L. and J.H. performed sequencing experiments. C.W. and P.G. performed sequencing experiments related to NMD inhibition. P.R.S., E.S., A.T.K., J.O., C.G.K., C.K., and U.N. contributed to study conception and experimental design. M.P. contributed conceptual input to the machine-learning component. S.M. supported biochemical assay setup. P.O. and A.R. supervised the project; contributed to experimental design across the study; provided resources; and co-wrote the manuscript; A.R. additionally contributed to the datascience work. All authors reviewed and approved the final manuscript.

## Supplementary Information

**Figure S1.**
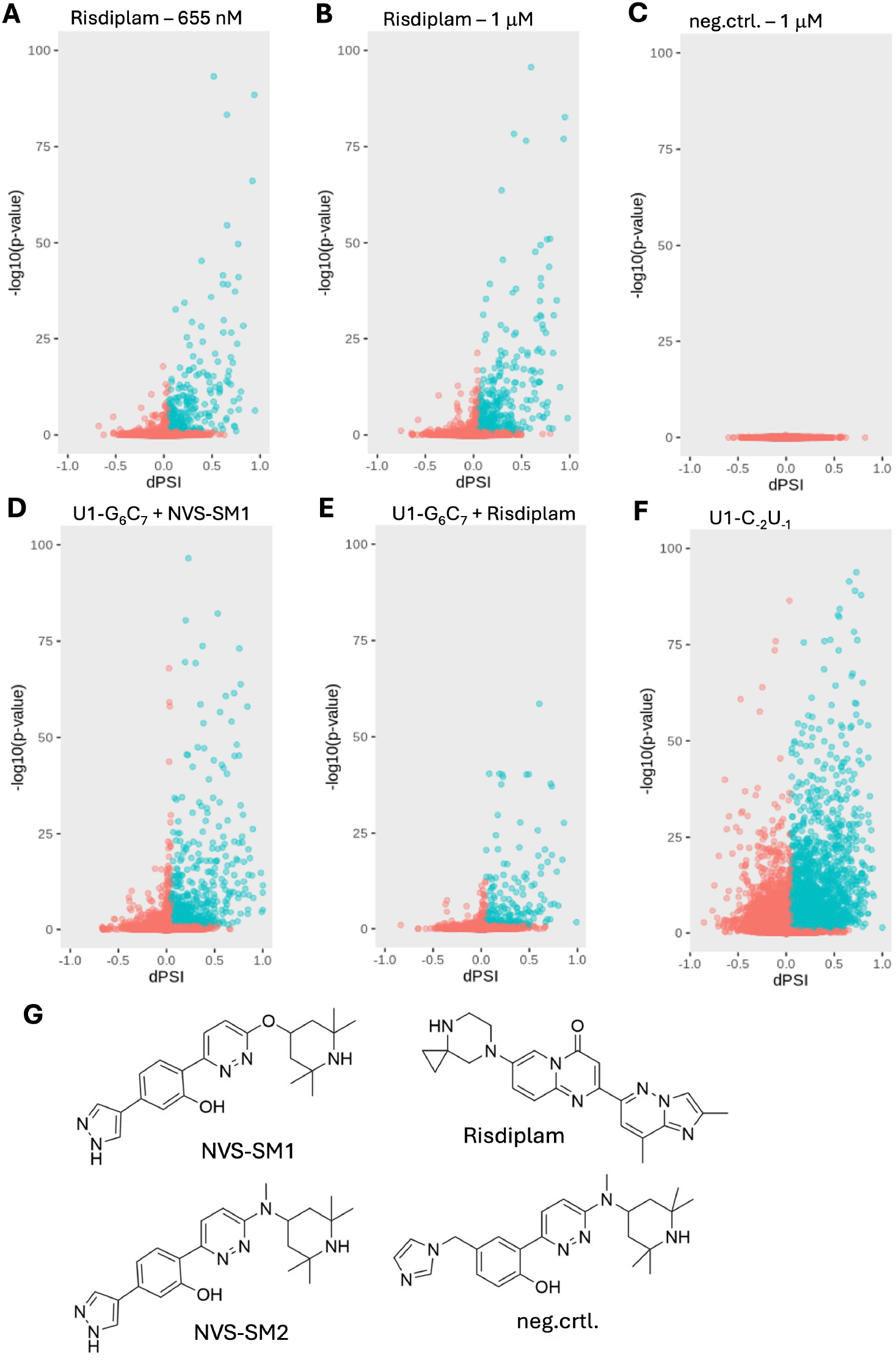
Differential splicing analysis between whole transcriptome profiles of SH-SY5Y cells treated with (A) 655 nM Risdiplam, (B) 1 µM Risdiplam or (C) 1 µM of inactive compound, and SH-SY5Y cells treated with DMSO. Differential splicing analysis between whole transcriptome profiles of HeLa cells transfected with U1-G_6_ C_7_ and treated with (D) 100 nM NVS-SM1 or (E) 1 µM Risdiplam, and transfected HeLa cells treated with DMSO. (F) Differential splicing analysis between whole transcriptome profiles of HeLa cells transfected with U1-C_−2_ U_−1_ and treated with DMSO and HeLa cells transfected with canonical U1. Exons highlighted in turquoise are considered as included in response to compound treatment (dPSI >0.05, adjusted p-value <0.1). (G) Chemical structures of the reference compounds used in this study: NVS-SM1, NVS-SM2, Risdiplam, and an inactive compound used as control (neg.ctrl.).

**Figure S2.**
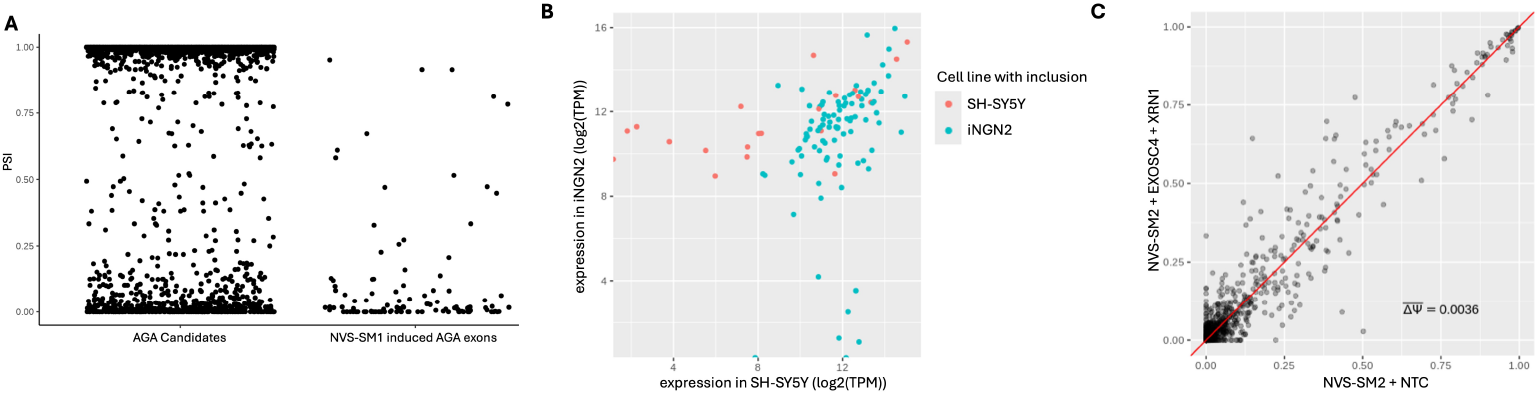
(A) Detected baseline inclusion in SH-SY5Y cells in the absence of a compound. Left group (AGA Candidates) represents all 5’ss matching an 5^*′*^-A_−3_ G_−2_ A_−1_ G_1_ U_2_ -3^*′*^ motif, identified as potential targets for NVS-SM1 in the human RNA-seq data collection from the Genotype-Tissue Expression Project. Right group (NVS-SM1 induced AGA exons) contains all AGA exons detected to be significantly induced in respond 25nM NVS-SM1 treatment. (B) Expression levels (log2(TPM)) of genes containing AGA-ending exons significantly induced only in SH-SY5Y cells or only in iNGN2 cells, respectivly. Expression in iNGN2 cells shown on y-axis and expression in SH-SY5Y cells shown on x-axis. (C) Scatter plot showing PSI values of AGA-ending exons in SH-SY5Y cells treated with 100nM of the NVS-SM2, an analog of NVS-SM1 (Supplementary Figure S1G), in the presence of NMD inhibiting siRNAs (EXOSC4 + XRN1) or non-targeting control (NTC). The red diagonal line indicates equal PSI values between the compared conditions and ΔΨ the mean change in inclusion.

**Figure S3.**
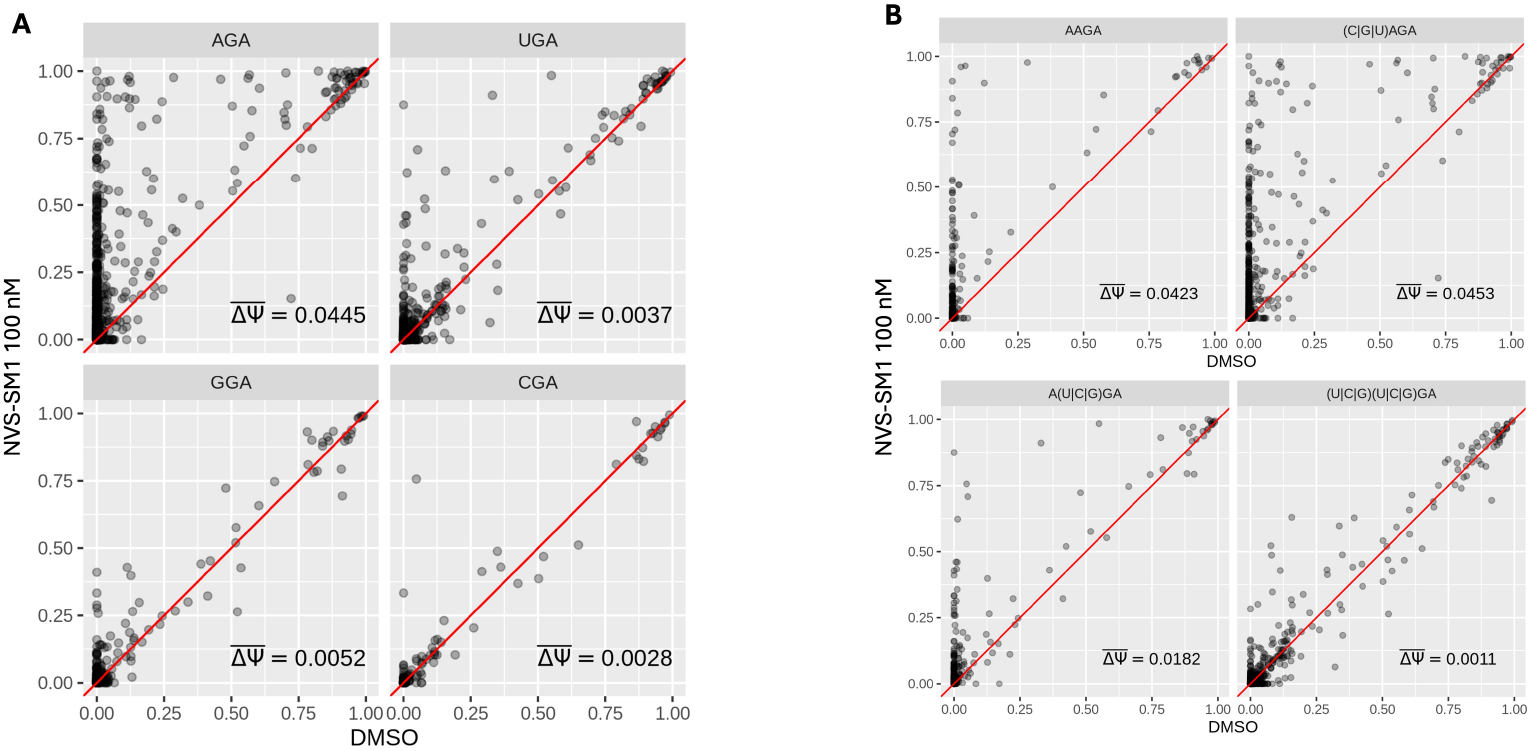
Scatter plots showing PSI values of GA-ending exons in cells treated with 100 nM NVS-SM1 or DMSO. (A) Exons grouped by their –3 position (AGA, UGA, GGA, or CGA). (B) Exons grouped according to adenine presence at the –3 and –4 positions: A at both positions (AAGA), A only at –3 ((C|G|U)AGA), A only at –4 (A(C|G|U)GA), or A absent at both ((C|G|U)(C|G|U)GA). The red diagonal line indicates equal PSI values between the compared conditions and ΔΨ the mean change in inclusion.

**Figure S4.**
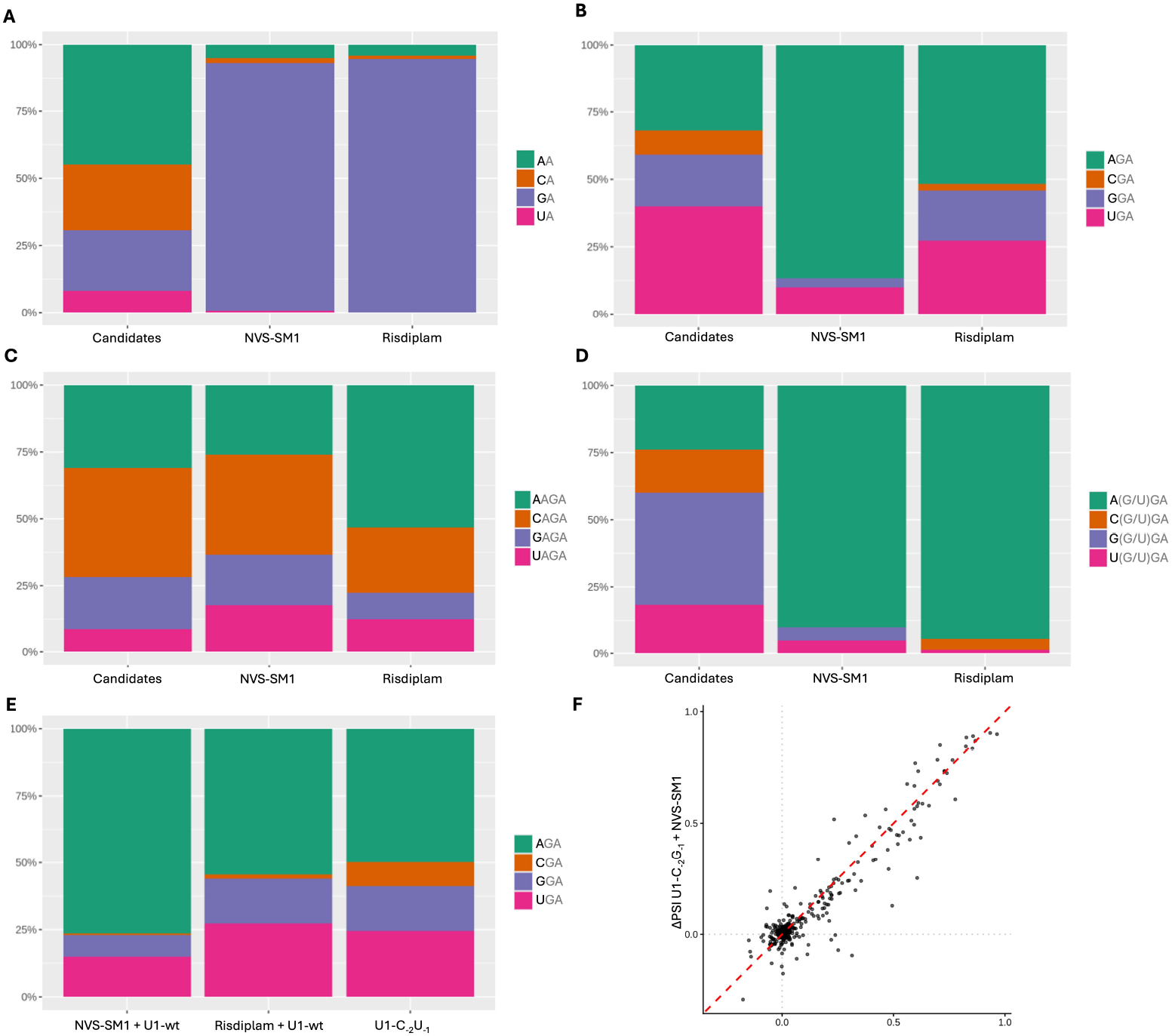
(A) Frequency of each nucleotide at position -2 within all A-ending exons (Candidates) or all induced A-ending exons upon NVS-SM1 or Risdiplam treatment. (B) Frequency of each nucleotide at position -3 within all GA-ending exons (Candidates) or all induced GA-ending exons upon NVS-SM1 or Risdiplam treatment in SH-SY5Y cells. (C) Frequency of each nucleotide at position -4 within all AGA exons (Candidates) or all induced AGA exons upon NVS-SM1 or Risdiplam treatment. (D) Frequency of each nucleotide at position -4 of all UGA or GGA exons (Candidates) or all induced UGA or GGA exons upon NVS-SM1 or Risdiplam treatment. (E) Frequency of each nucleotide at position -3 of GA-ending exons induced in response to NVS-SM1 or Risdiplam treatment in HeLa cells transfected with wt U1 snRNA or in cells transfected with U1-C_−2_ U_−1_ and treated with DMSO.

**Figure S5.**
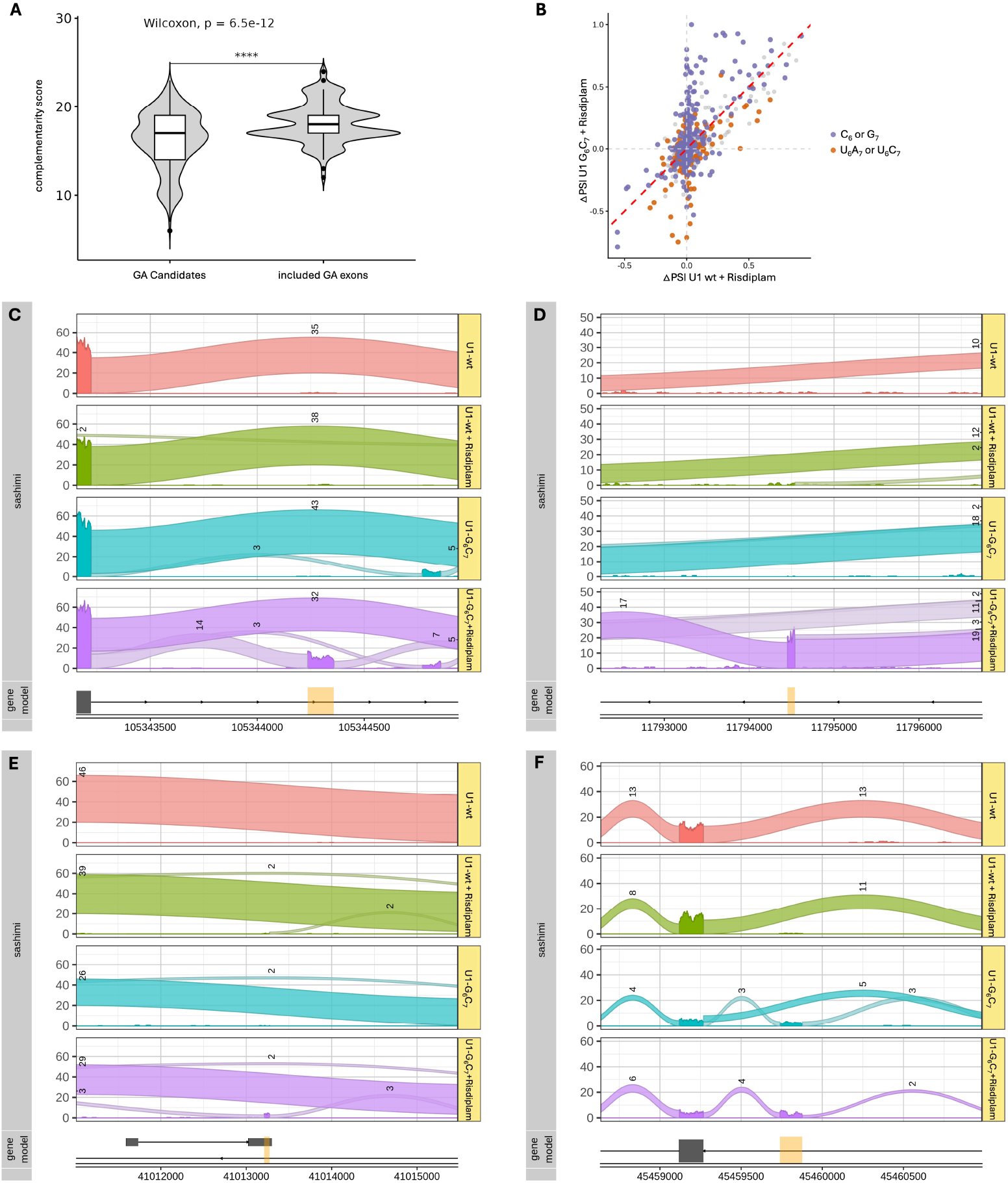
(A) Violin plot showing the complementarity between the U1 snRNA and U1 binding sites of GA-ending exons included in response to Risdiplam (included GA exons) and those GA exons that were potential targets for Risdiplam (GA Candidates) (p-value < 0.0001). The complementarity score was calculated by summing up all H-bonds possibly formed between the two RNAs. (B) Scatter plot comparing exon inclusion changes (ΔPSI) upon compound treatment in cells expressing canonical (U1-wt + Risdiplam) or mutated U1 snRNA (U1-G_6_ C_7_ + Risdiplam). The x-axis shows values for exons in cells transfected with canonical U1 and treated with 100 nM Risdiplam, and the y-axis shows values for the same exons in cells transfected with U1-G_6_ C_7_ under identical treatment conditions. ΔPSI between the two conditions. Exons with higher complementarity to the mutated U1 (carrying a C at position 6 or a G at position 7 of the U1 binding site) are highlighted in purple, whereas exons with higher complementarity to the canonical U1 are shown in orange. The diagonal line indicates equal ΔPSI between the two conditions. (C) Sashimi plots of RNA-seq results deriving from the same experiment as explained in Figure 3A. Plot highlights an exon within the gene ERGIC1. Significant inclusion only detected in cells transfected with U1-G_6_ C_7_ and treated with 1µM Risdiplam (dPSI = 0.36, p-value < 0.001). (D) Sashimi plots of RNA-seq results deriving from the same experiment as explained in Figure 3A. Plot highlights an exon within the gene *TAMM41*. Significant inclusion only detected in cells transfected with U1-G_6_ C_7_ and treated with 1µM Risdiplam (dPSI = 0.73, p-value < 0.001). (E) Sashimi plots of RNA-seq results deriving from the same experiment as explained in Figure 3A. Plot highlights an exon within the gene *INO80*. Low, but not significant inclusion detected in cells transfected with canonical U1 snRNA and 1µM Risdiplam. Significant inclusion only detected in cells transfected with U1-G_6_ C_7_ and Risdiplam treatment (dPSI = 0.17, p-value < 0.001). (F) Sashimi plots of RNA-seq results deriving from the same experiment as explained in Figure 3A. Plot highlights an exon within the gene MARCHF8. Significant inclusion detected in cells transfected with U1-G_6_ C_7_ (dPSI = 0.79, p-value < 0.001) and boosted inclusion in cells transfected with U1-G_6_ C_7_ and additional Risdiplam treatment (dPSI = 1.0, p-value < 0.001).

**Figure S6.**
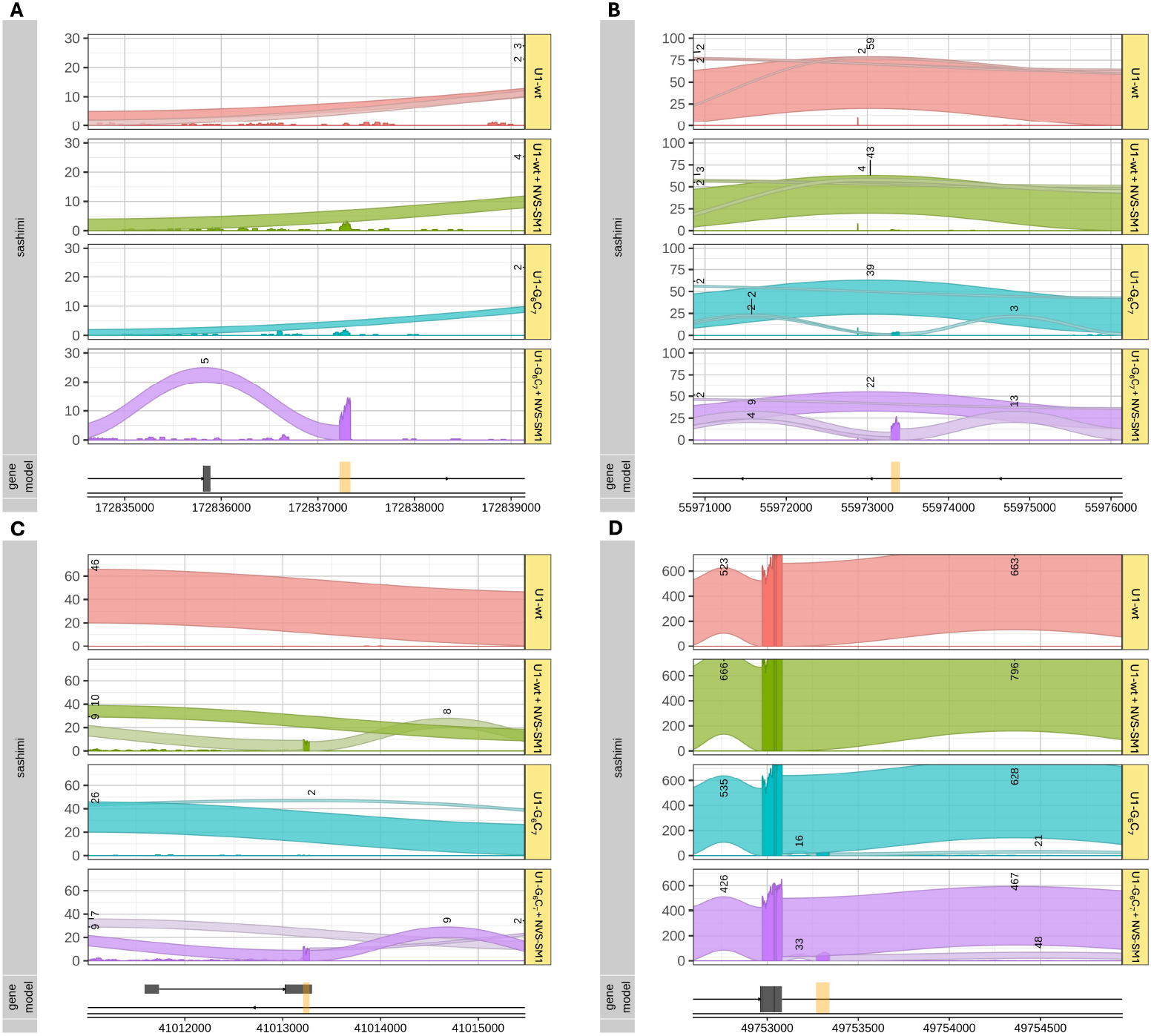
(A) Sashimi plots of RNA-seq results deriving from the same experiment as in Figure 3A. Plot highlights an exon within the gene *ERGIC1*. Significant inclusion only detected in cells transfected with U1-G_6_ C_7_ and treated with 100nM NVS-SM1 dPSI = 0.13, p-value < 0.001). (B) Sashimi plots of RNA-seq results deriving from the same experiment as explained in Figure 3A. Plot highlights an exon within the gene *IL6ST*. Significant inclusion detected in cells transfected with U1-G_6_ C_7_ (dPSI = 0.1, p-value < 0.001) and boosted inclusion in cells additionally treated with 100nM NVS-SM1 (dPSI = 0.43, p-value < 0.001). (C) Sashimi plots of RNA-seq results deriving from the same experiment as explained in Figure 3A. Plot highlights an exon within the gene *INO80*. Significant inclusion detected in cells transfected with canonical U1 snRNA (U1-wt) and NVS-SM1 treatment (dPSI = 0.6, p-value < 0.001) and boosted inclusion in cells transfected with U1-G_6_ C_7_ and NVS-SM1 treatment (dPSI = 0.7, p-value < 0.001). (D) Sashimi plots of RNA-seq results deriving from the same experiment as explained in Figure 3A. Plot highlights an exon within the gene *TMBIM6*. Significant inclusion detected in cells transfected with U1-G_6_ C_7_ (dPSI = 0.06, p-value < 0.001) and boosted inclusion in cells additionally treated with 100nM NVS-SM1 (dPSI = 0.12, p-value < 0.001).

**Figure S7.**
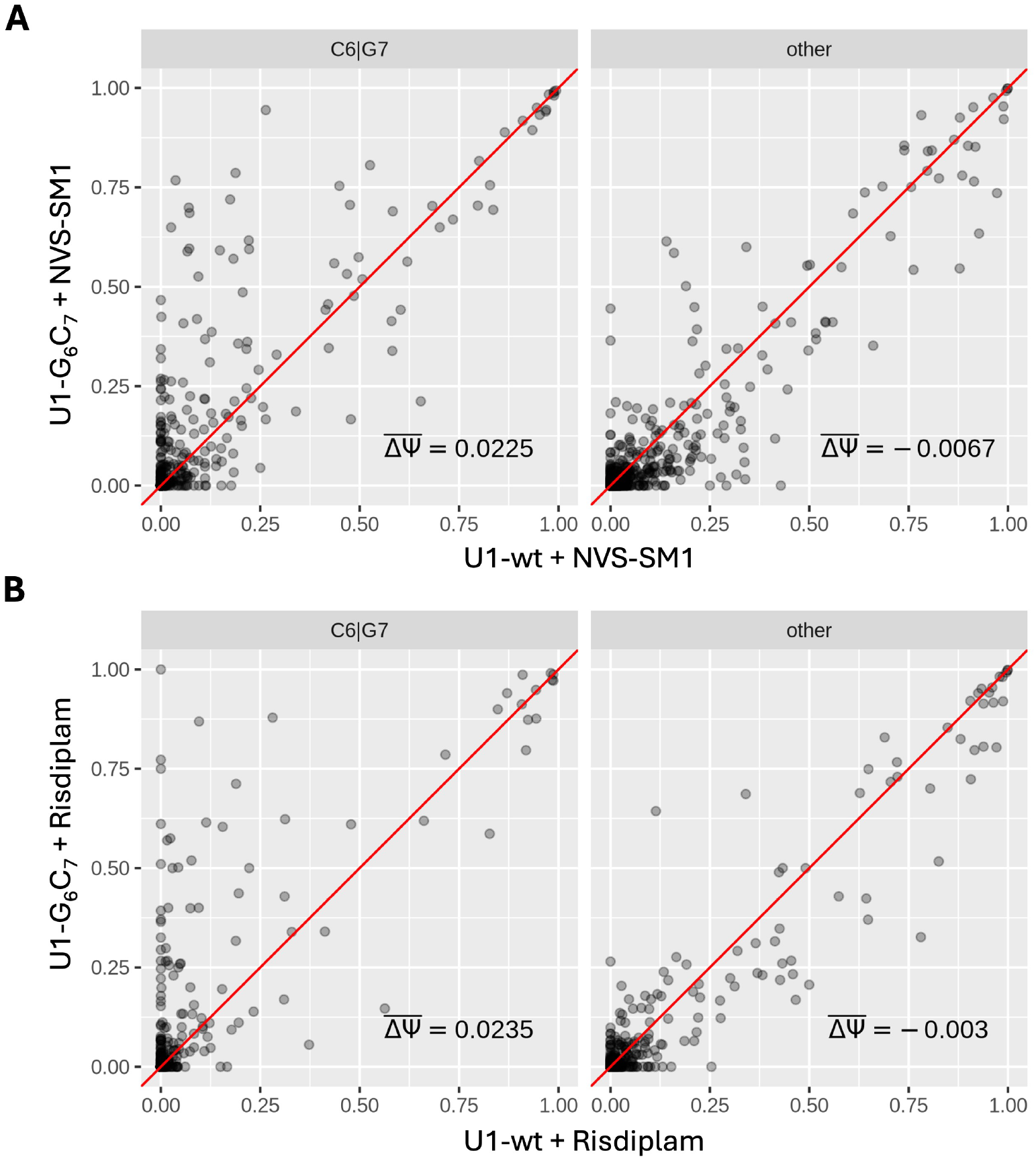
Scatter plots showing PSI values of GA-ending exons, grouped by the presence of either a C at position +6 or a G at +7 (C6|G7) versus exons lacking one of these nucleotides (other). (A) Cells treated with NVS-SM1 and transfected with either U1-G_6_ C_7_ or canonical U1 snRNA. (B) Cells treated with Risdiplam and transfected with either U1-G_6_ C_7_ or canonical U1 snRNA. The red diagonal line indicates equal PSI values between the compared conditions and ΔΨ the mean change in inclusion.

**Figure S8.**
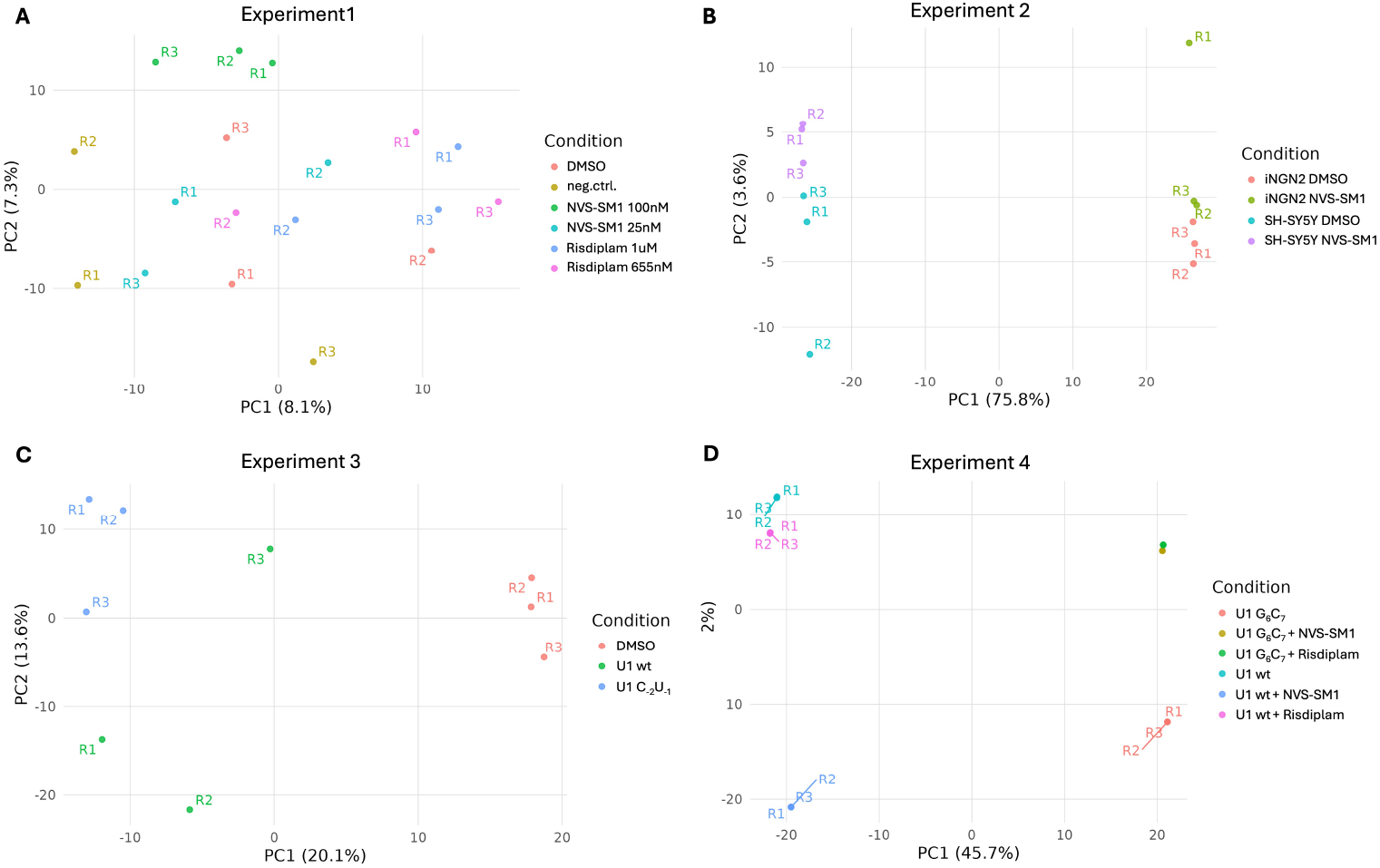
Principal component analysis (PCA) of exon inclusion profiles. PCA was performed on the 1,000 most variable exons across samples. (A) SH-SY5Y cells treated with DMSO, negative control compound, Risdiplam, or NVS-SM1 at different concentrations, each in biological triplicates (R1–R3). (B) Comparison of compound-treated and untreated samples in different cell types (SH-SY5Y and iNGN2). (C) HEK293 cells expressing U1-wt or U1-C_−2_ U_−1_ snRNA, with and without compound treatment. (D) HEK293 cells expressing either canonical U1 or mutated U1 snRNA (U1-G_6_ C_7_), with and without compound treatment. Each point represents one replicate, and proximity indicates similarity in exon inclusion patterns.

**Figure S9.**
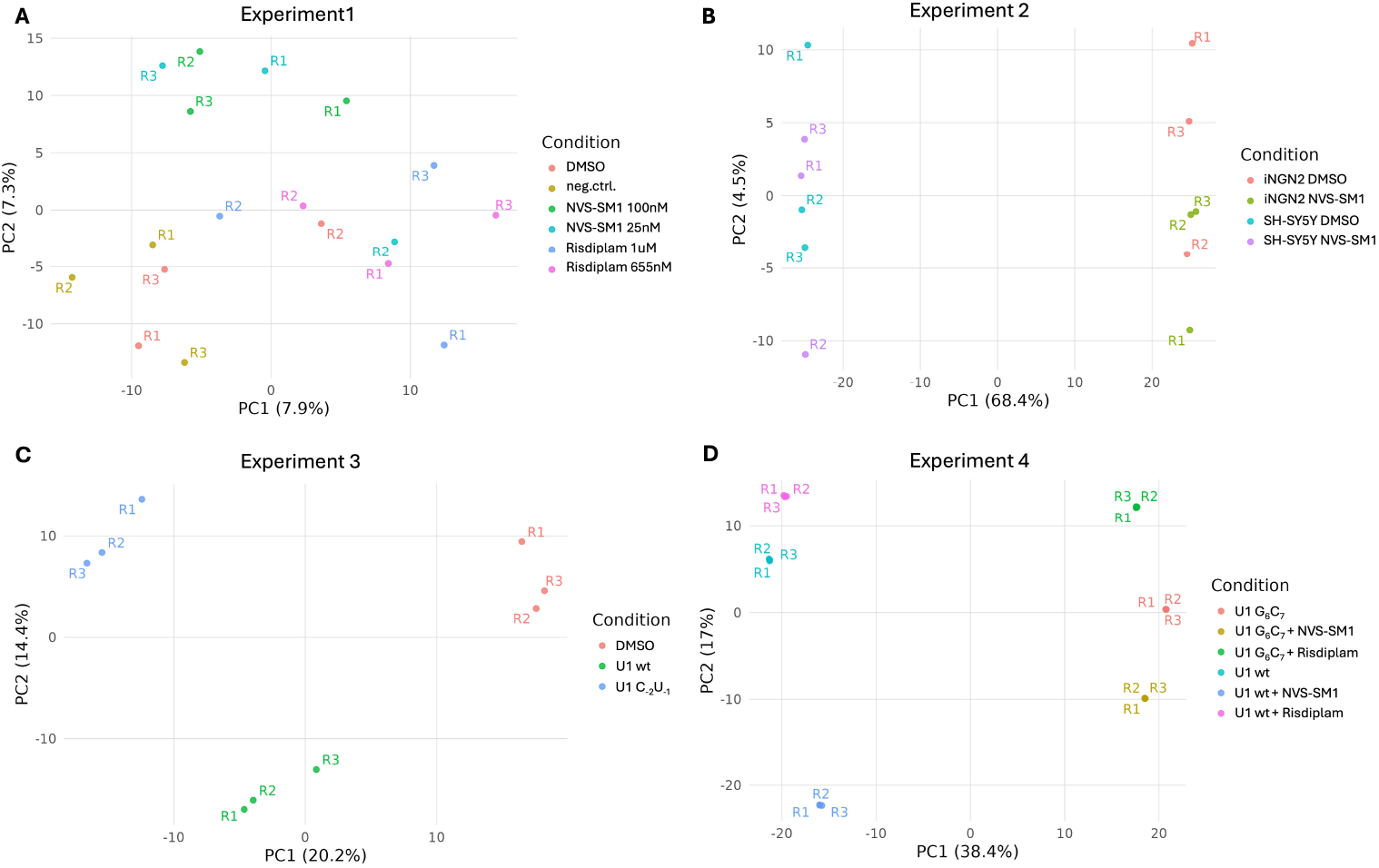
PCA of GA-ending exons. (A) SH-SY5Y cells treated with DMSO, negative control compound, Risdiplam, or NVS-SM1 at different concentrations, each in biological triplicates (R1–R3). (B) Comparison of compound-treated and untreated samples in different cell types (SH-SY5Y and iNGN2). (C) HEK293 cells expressing U1-wt or U1-C_−2_ U_−1_ snRNA, with and without compound treatment. (D) HEK293 cells expressing either canonical U1 or mutated U1 snRNA, with and without compound treatment. Each dot represents one replicate; clustering reflects similarity in exon inclusion profiles among conditions.

**Figure S10.**
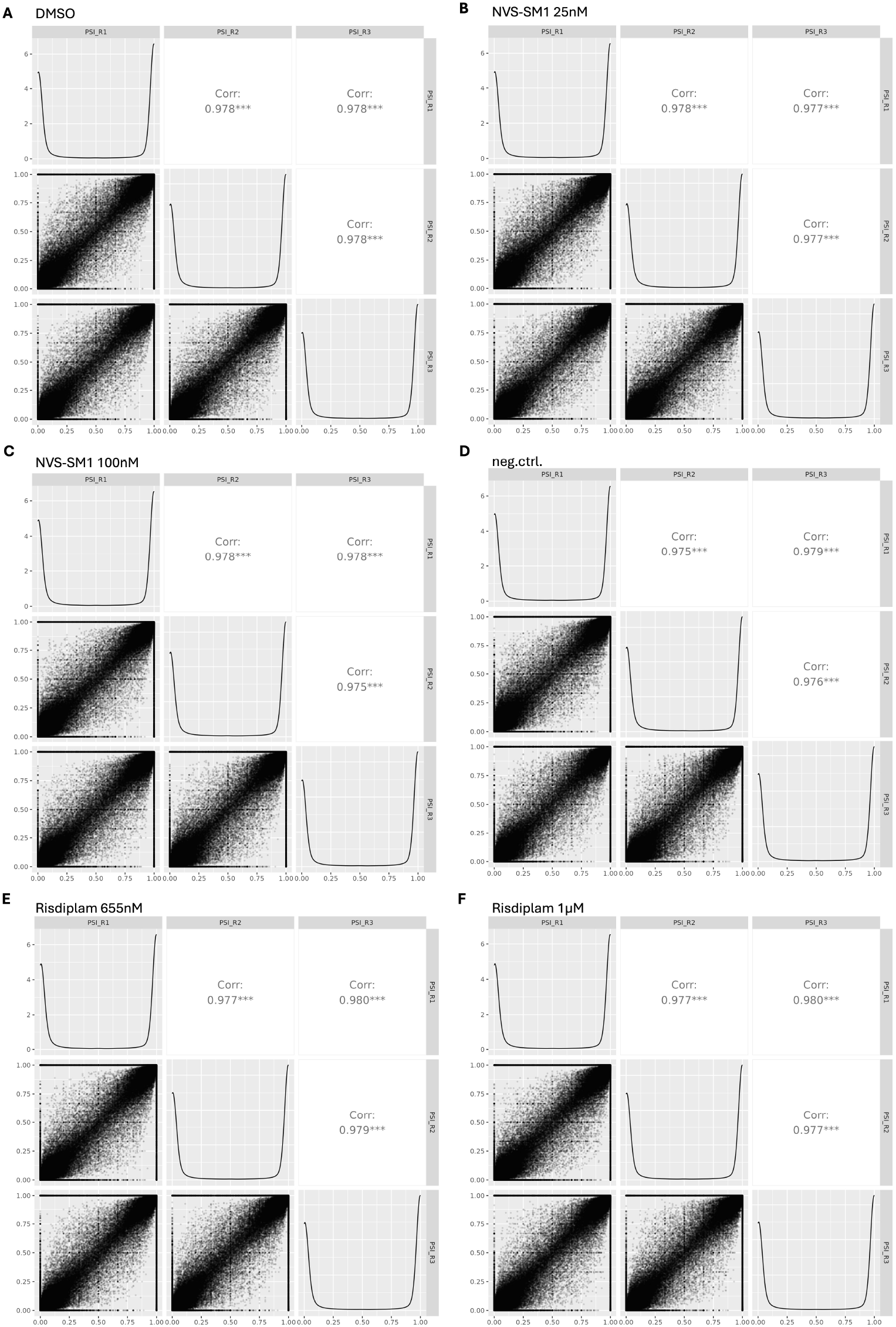
Reproducibility of exon inclusion measurements across biological replicates. Pairwise correlations of PSI values between triplicates for SH-SY5Y cells treated with (A) DMSO, (B) 25 nM NVS-SM1, (C) 100 nM NVS-SM1, (D) negative control compound, (E) 655 nM Risdiplam, and (F) 1 µM Risdiplam. Each panel shows pairwise scatter plots of PSI values between replicates (R1–R3), with Pearson correlation coefficients indicated. High correlations across all conditions demonstrate the reproducibility of exon inclusion quantification.

**Figure S11.**
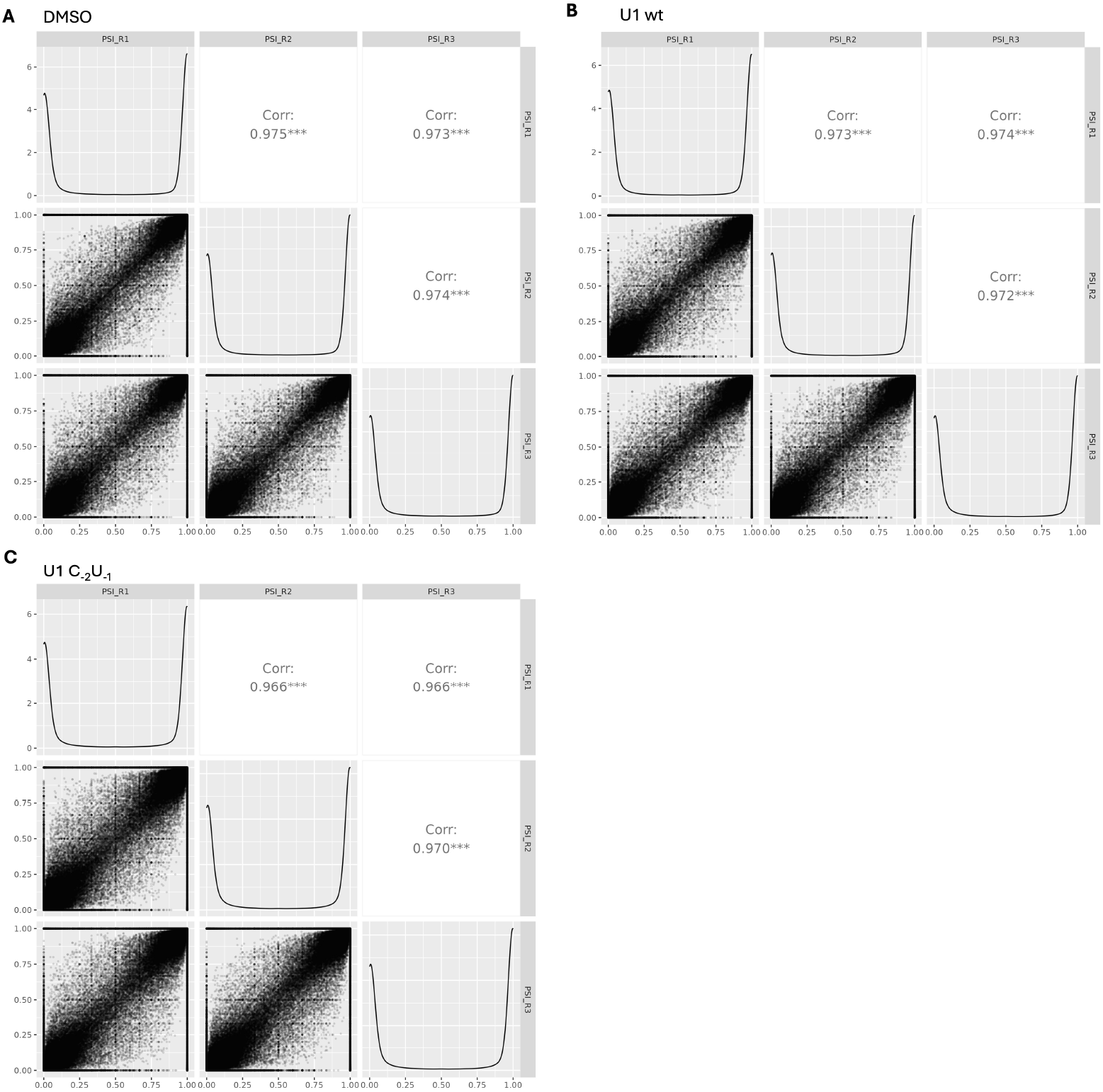
Reproducibility of exon inclusion measurements across biological replicates in U1-transfected cells. Pairwise correlations of PSI values between replicates for cells transfected with (A) DMSO control, (B) canonical U1 snRNA (U1 wt), and (C) mutated U1 snRNA (U1-C_−2_ U_−1_). Each panel shows pairwise scatter plots of PSI values between replicates (R1–R3), with Pearson correlation coefficients indicated. High correlation coefficients across all samples confirm the reproducibility of exon inclusion quantification.

**Figure S12.**
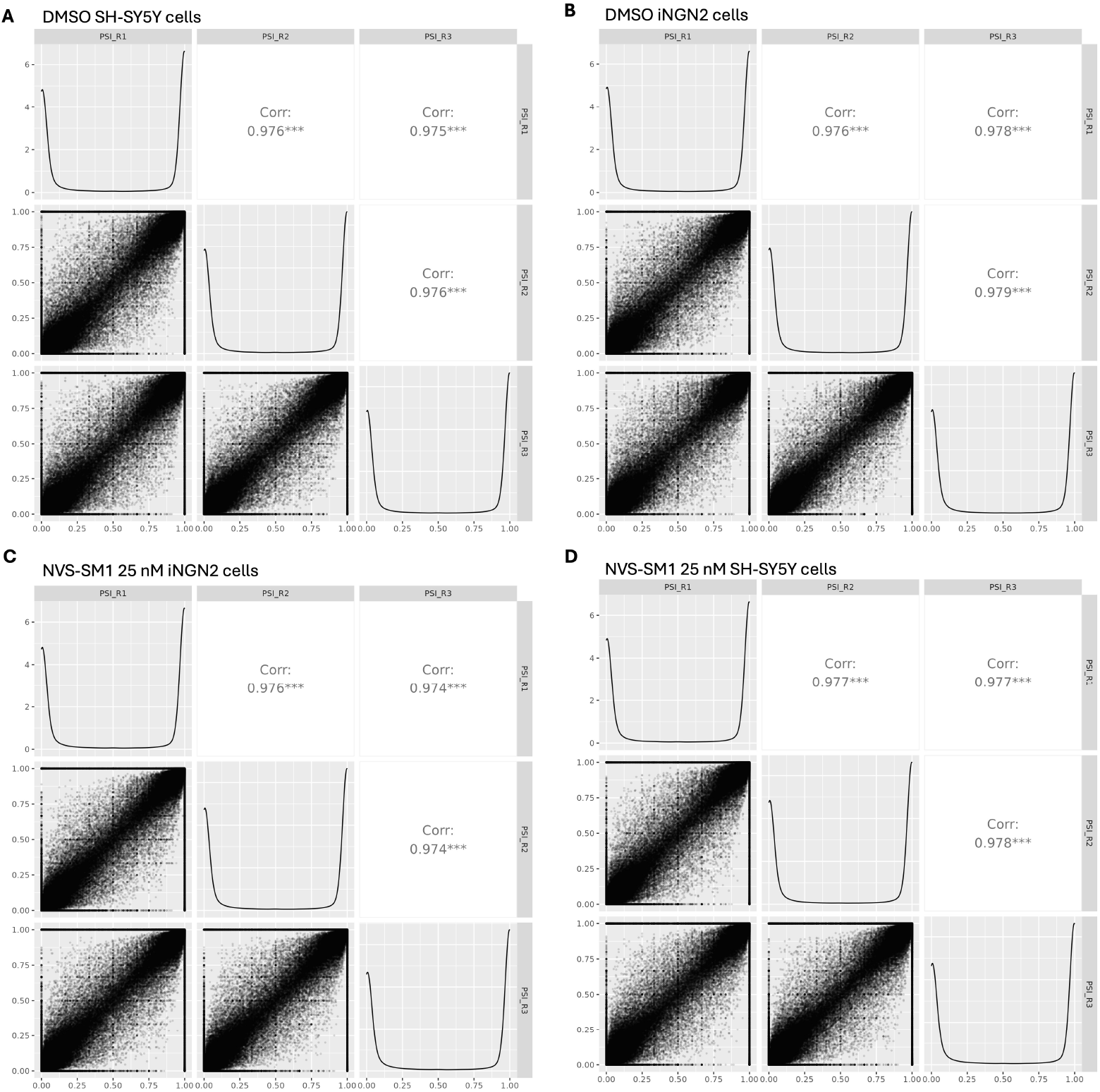
Reproducibility of exon inclusion measurements across biological replicates in SH-S5Y5 and iNGN2 cells. Pairwise correlations of PSI values between replicates for SH-S5Y5 and iNGN2 cells treated with either DMSO (A and B) or 25 nM NVS-SM1 (C and D). Each panel shows pairwise scatter plots of PSI values between replicates (R1–R3), with Pearson correlation coefficients displayed. High correlations across all conditions confirm the robustness and reproducibility of exon inclusion quantification.

**Figure S13.**
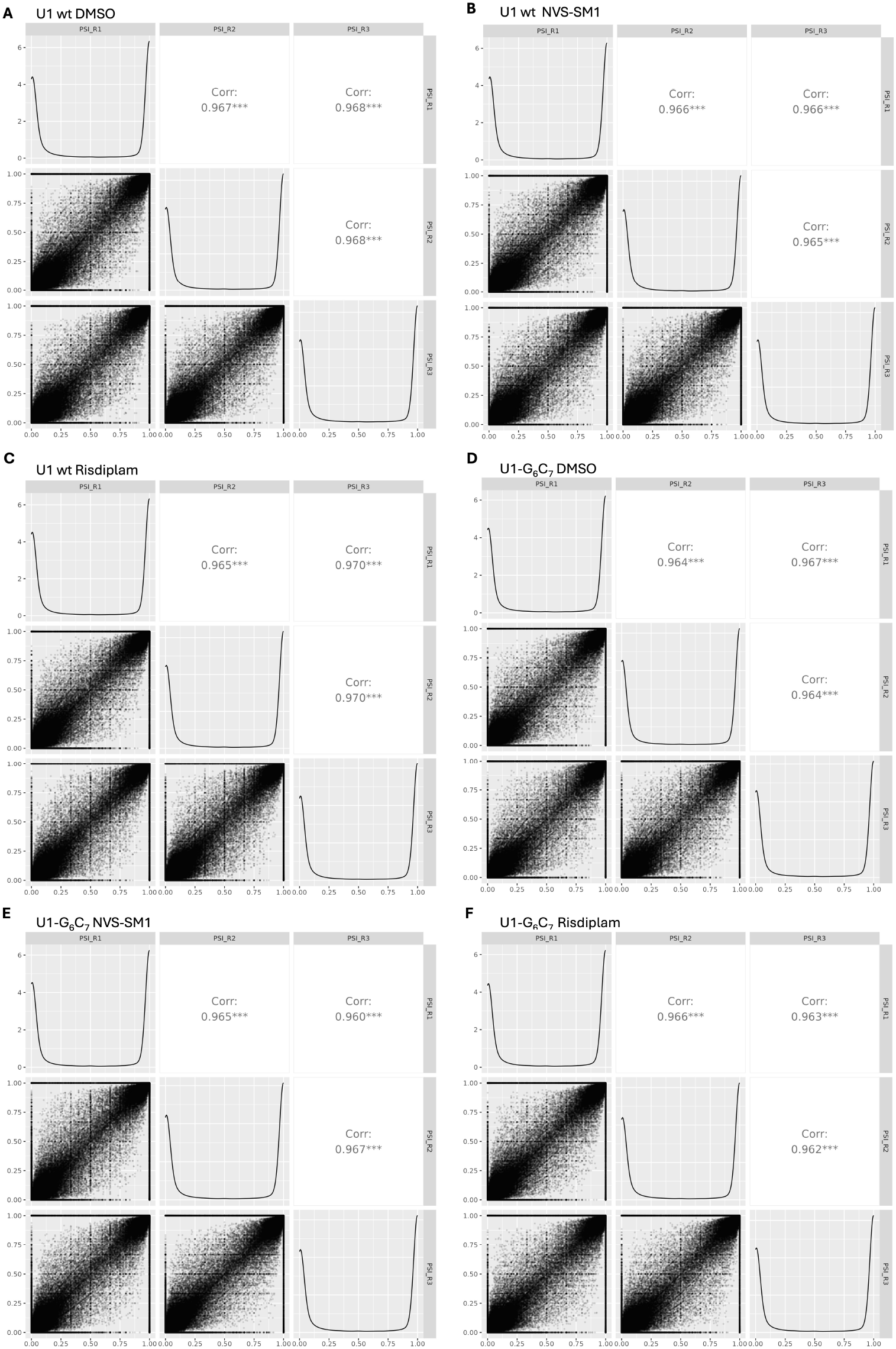
Reproducibility of exon inclusion measurements across biological replicates in HEK293 cells expressing canonical or mutant U1 snRNA. Pairwise correlations of PSI values between replicates for cells expressing either wild-type U1 (A–C) or U1-G_6_ C_7_ (D–F), treated with DMSO, NVS-SM1, or Risdiplam as indicated. Each panel shows pairwise scatter plots of PSI values between replicates (R1–R3), with Pearson correlation coefficients displayed. High correlations across all conditions confirm the robustness and reproducibility of exon inclusion quantification.

**Figure S14.**
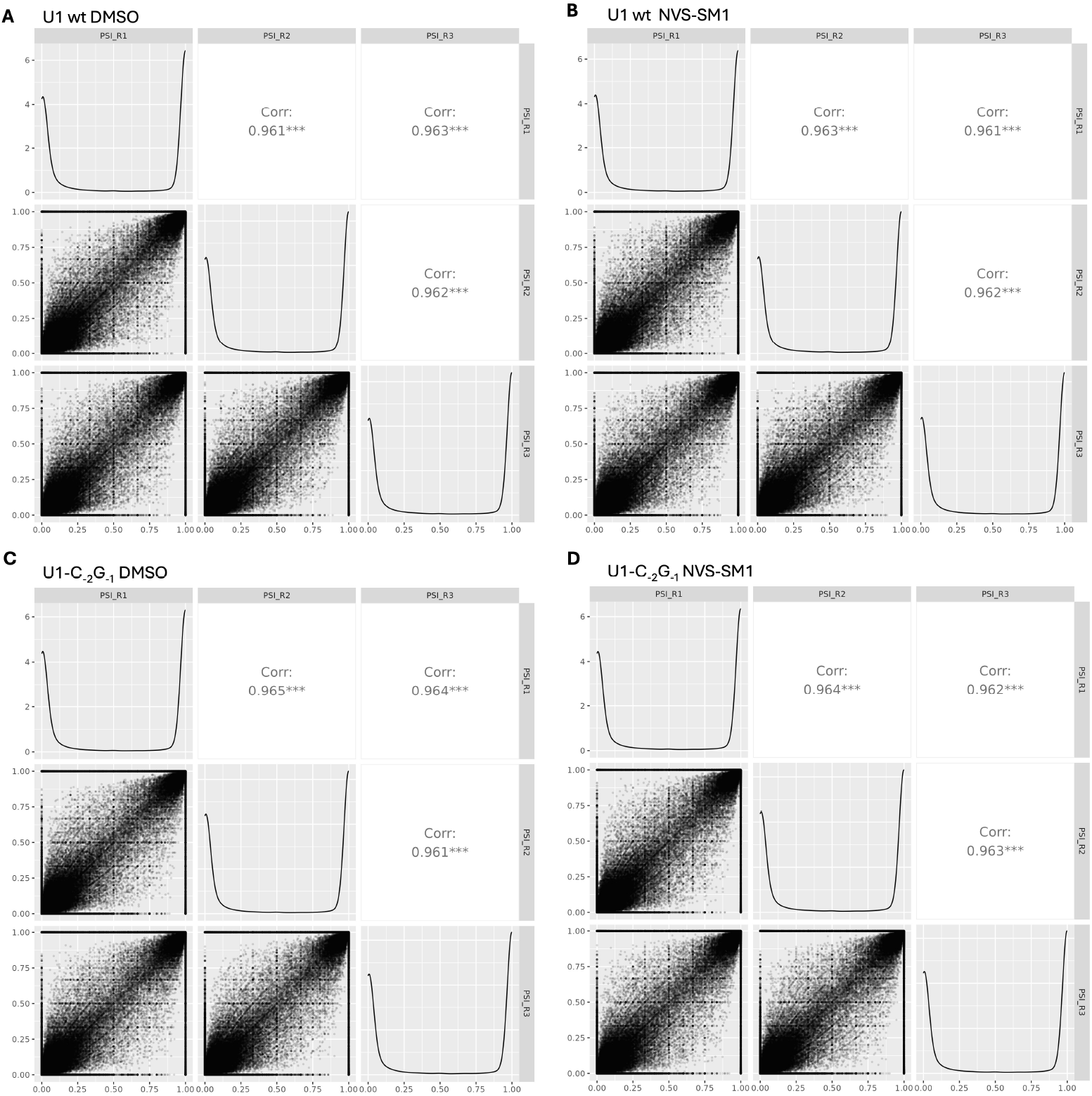
Reproducibility of exon inclusion measurements across biological replicates in SH-SYS5 cells expressing canonical or mutant U1 snRNA. Pairwise correlations of PSI values between replicates for cells expressing either wild-type U1 (A–B) or U1-C_−2_ G_−1_ (C–D), treated with DMSO or NVS-SM1 as indicated. Each panel shows pairwise scatter plots of PSI values between replicates (R1–R3), with Pearson correlation coefficients displayed. High correlations across all conditions confirm the robustness and reproducibility of exon inclusion quantification.

**Figure S15.**
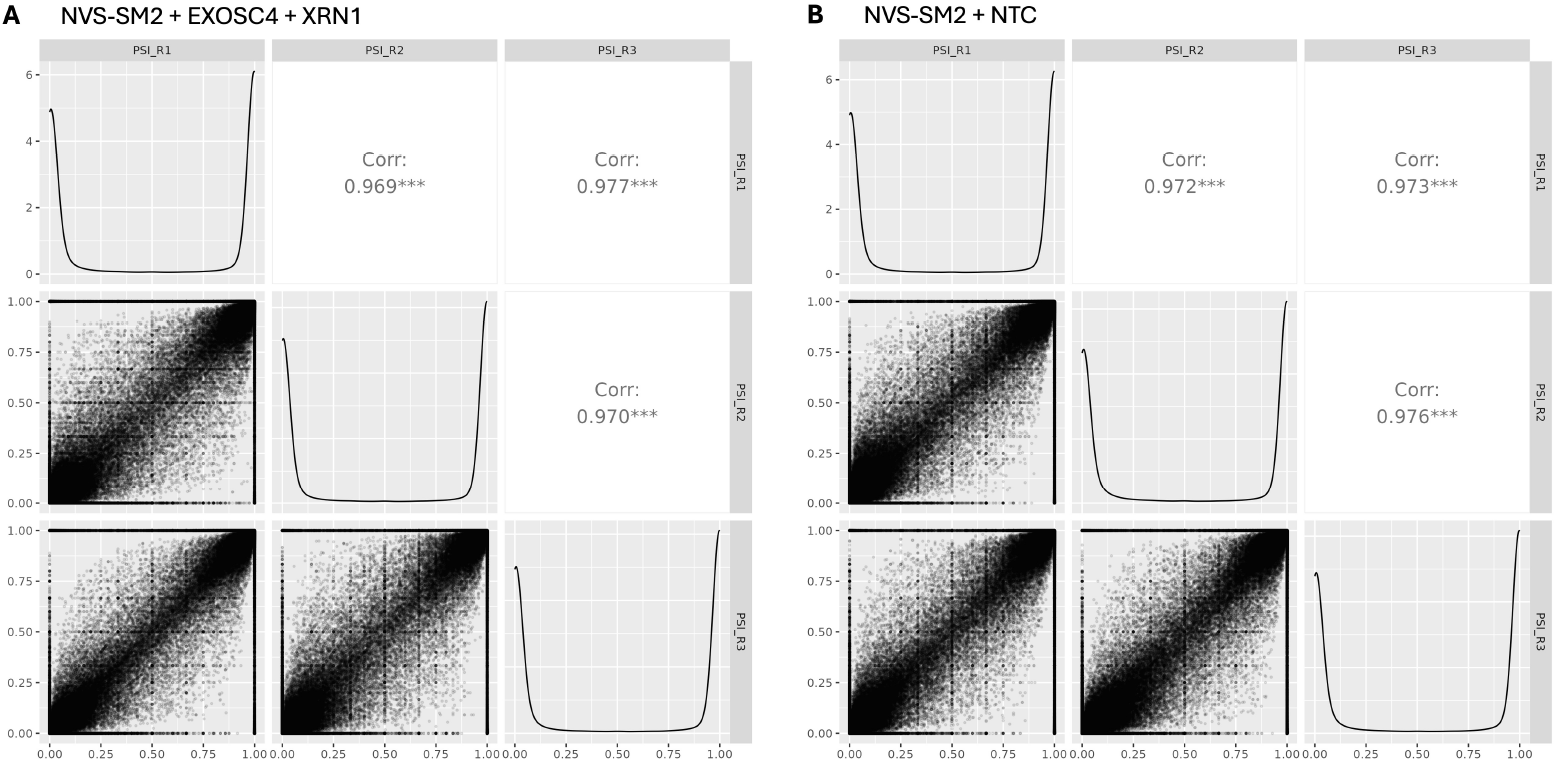
Reproducibility of exon inclusion measurements across biological replicates in SH-SYS5 cells expressing canonical or mutant U1 snRNA. Pairwise correlations of PSI values between replicates for cells with XRN1 and EXOSC4 knockdown (A) or transfected with non-targeting control (B), treated with 100 nM of NVS-SM2. Each panel shows pairwise scatter plots of PSI values between replicates (R1–R3), with Pearson correlation coefficients displayed. High correlations across all conditions confirm the robustness and reproducibility of exon inclusion quantification.

**Table S1.** List of some NVS-SM1-induced exons with corresponding complementarity scores. Scores were calculated by summing the maximum number of possible H-bonds formed between the 5’ss of AGA exons and U1 snRNA. We also considered the possibility of an A_−1_ bulge formation, thus including the -4 position of the exon in the score calculation.

| Gene name | chr. | start | end | U1 binding site | Complementarity score |
| --- | --- | --- | --- | --- | --- |
| *HTT* | 4 | 3213622 | 3213736 | CAGAGUAAGGGG | 20 |
| *KCNT2* | 1 | 196317148 | 196317173 | CAGAGUAAGAAA | 21 |
| *LRRC42* | 1 | 53947862 | 53947981 | UAGAGUAAGUUA | 19 |
| *MLLT10* | 10 | 21728632 | 21728675 | GAGAGUAAGAGA | 18 |
| *ARHGAP1* | 11 | 46697021 | 46697069 | CAGAGUAAGAGU | 20 |
| *WNK1* | 12 | 895173 | 895196 | GAGAGUAGGUGA | 19 |

## Bibliography

Bhattacharyya, A. et al. Small molecule splicing modifiers with systemic htt-lowering activity. Nature Communications, 12(1):7299, 2021. doi: 10.1038/s41467-021-27661-5.

Black, D. L. Mechanisms of alternative pre-messenger rna splicing. Annual Review of Biochemistry, 72:291–336, 2003. doi: 10.1146/annurev.biochem.72.121801.161720.

Borowsky, B., Ramos, H., Caputo, A., Hartmann, A., Faller, T., Peters, T., Sui, Y., Liu, F., Meadowcroft, M., David, O. J., Laisney, M., Kinhikar, A., Marder, K. S., Tabrizi, S. J., Landwehrmeyer, G. B., and Leavitt, B. R. Oral splicing modulator branaplam in huntington’s disease: a phase 2 randomized controlled trial. Nature Medicine, 32(1):103–112, 2026. doi: 10.1038/s41591-025-04117-4.

Bradrick, T. D. and Marino, J. P. Ligand-induced changes in 2-aminopurine fluorescence as a probe for small molecule binding to hiv-1 tar rna. RNA, 10(9):1459–1468, 2004. doi: 10.1261/rna.7650704.

Campagne, S. et al. Structural basis of a small molecule targeting rna for a specific spliing correction. Nature Chemical Biology, 15(12):1191–1198, 2019. doi: 10.1038/s41589-019-0377-2.

Campagne, S., Allain, F. H.-T., and colleagues. The diversity of splicing modifiers acting on a-1 bulged 5 splice sites reveals rules for rational drug design. Nucleic Acids Research, 52(X):XXXX–XXXX, 2024. doi: 10.1093/nar/xxxxxx.

Cartegni, L., Wang, J., Zhu, Z., Zhang, M. Q., and Krainer, A. R. ESEfinder: A web resource to identify exonic splicing enhancers. Nucleic Acids Research, 31(13):3568–3571, 2003. doi: 10.1093/nar/gkg616.

Chen, J. L. et al. Design, optimization, and study of small molecules that target tau pre-mrna and affect splicing. Journal of the American Chemical Society, 142(19):8706–8727, 2020. doi: 10.1021/jacs.0c02213.

Dhillon, S. Risdiplam: First approval. Drugs, 80(17):1853–1858, 2020. doi: 10.1007/s40265-020-01394-7.

Freund, M. et al. Extended base pair complementarity between u1 snrna and the 5’ splice site does not inhibit splicing in higher eukaryotes, but rather increases 5’ splice site recognition. Nucleic Acids Research, 33(16):5112–5119, 2005. doi: 10.1093/nar/gki812.

Havens, M. A. and Hastings, M. L. Splice-switching antisense oligonucleotides as therapeutic drugs. Nucleic Acids Research, 44(14):6549–6563, 2016. doi: 10.1093/nar/gkw589.

Ishigami, Y. et al. Specificity, synergy, and mechanisms of splice-modifying drugs. Nature Communications, 15(1), 2024. doi: 10.1038/s41467-024-46090-5.

Keller, C. G. et al. An orally available, brain penetrant, small molecule lowers huntingtin levels by enhancing pseudoexon inclusion. Nature Communications, 13(1):1150, 2022. doi: 10.1038/s41467-022-28847-4.

Krach, F. et al. An alternative splicing modulator decreases mutant htt and improves the molecular fingerprint in huntington’s disease patient neurons. Nature Communications, 13(1):6797, 2022. doi: 10.1038/s41467-022-34396-0.

Le, T. T. et al. Smndelta7, the major product of the centromeric survival motor neuron (smn2) gene, extends survival in mice with spinal muscular atrophy and associates with full-length smn. Human Molecular Genetics, 14(6):845–857, 2005. doi: 10.1093/hmg/ddi078.

Lee, Y. and Rio, D. C. Mechanisms and regulation of alternative pre-mrna splicing. Annual Review of Biochemistry, 84:291–323, 2015. doi: 10.1146/annurev-biochem-060614-034316.

Lefebvre, S. et al. Identification and characterization of a spinal muscular atrophy-determining gene. Cell, 80(1):155–165, 1995. doi: 10.1016/0092-8674(95)90460-3.

Liu, H. X., Zhang, M., and Krainer, A. R. Identification of functional exonic splicing enhancer motifs recognized by SF2/ASF and SC35. Proceedings of the National Academy of Sciences, 95(18):10626–10631, 1998. doi: 10.1073/pnas.95.18.10626.

Malard, F. et al. The diversity of splicing modifiers acting on a-1 bulged 5’-splice sites reveals rules for rational drug design. Nucleic Acids Research, 52(8):4124–4136, 2024. doi: 10.1093/nar/gkae200.

Mendell, J. R. et al. Eteplirsen for the treatment of duchenne muscular dystrophy. Annals of Neurology, 74(5):637–647, 2013. doi: 10.1002/ana.23982.

Mercuri, E. et al. Nusinersen versus sham control in later-onset spinal muscular atrophy. New England Journal of Medicine, 378(7):625–635, 2018. doi: 10.1056/NEJMoa1710504.

Monani, U. R. et al. A single nucleotide difference that alters splicing patterns distinguishes the sma gene smn1 from the copy gene smn2. Human Molecular Genetics, 8(7):1177– 1183, 1999. doi: 10.1093/hmg/8.7.1177.

Palacino, J. et al. Smn2 splice modulators enhance u1-pre-mrna association and rescue sma mice. Nature Chemical Biology, 11(7):511–517, 2015. doi: 10.1038/nchembio.1834.

Papasaikas, P. and Valcarcel, J. The spliceosome: The ultimate rna chaperone and sculptor. Trends in Biochemical Sciences, 41(1):33–45, 2016. doi: 10.1016/j.tibs.2015.11.003.

Rachofsky, E. L., Osman, R., and Ross, J. B. Probing structure and dynamics of dna with 2-aminopurine: Effects of local environment on fluorescence. Biochemistry, 40(4):946– 956, 2001. doi: 10.1021/bi001763s.

Ratni, H. et al. Specific correction of alternative survival motor neuron 2 splicing by small molecules: Discovery of a potential novel medicine to treat spinal muscular atrophy. Journal of Medicinal Chemistry, 59(13):6086–6100, 2016. doi: 10.1021/acs.jmedchem.6b00474.

Ratni, H. et al. Discovery of risdiplam, a selective survival of motor neuron-2 (smn2) gene splicing modifier for the treatment of spinal muscular atrophy (sma). Journal of Medicinal Chemistry, 61(15):6501–6517, 2018. doi: 10.1021/acs.jmedchem.8b00741.

Rau, M. J. and Hall, K. B. 2-aminopurine fluorescence as a probe of local rna structure and dynamics and global folding. Methods in Enzymology, 558:99–124, 2015. doi: 10.1016/bs.mie.2015.02.005.

Reyes, A. and Huber, W. Alternative start and termination sites of transcription drive most transcript isoform differences across human tissues. Nucleic Acids Research, 46(2):582– 592, 2018. doi: 10.1093/nar/gkx1165.

Ruggiu, M. et al. A role for smn exon 7 splicing in the selective vulnerability of motor neurons in spinal muscular atrophy. Molecular and Cellular Biology, 32(1):126–138, 2012. doi: 10.1128/MCB.06077-11.

Schafer, S. et al. Alternative splicing signatures in rna-seq data: Percent spliced in (psi). Current Protocols in Human Genetics, 87:11.16.1–11.16.14, 2015. doi: 10.1002/0471142905.hg1116s87.

Singh, R. N., Seo, J., and Singh, N. N. Rna in spinal muscular atrophy: Therapeutic implications of targeting. Expert Opinion on Therapeutic Targets, 24(8):731–743, 2020. doi: 10.1080/14728222.2020.1799523.

Tang, Z. et al. Rna-targeting splicing modifiers: Drug development and screening assays. Molecules, 26(8), 2021. doi: 10.3390/molecules26082253.

Tang, Q., Rodriguez, J., and Spitale, R. C. Recognition of single-stranded nucleic acids by small-molecule splicing modulators. Nucleic Acids Research, 49(14):7870–7883, 2021. doi: 10.1093/nar/gkab558.

Wang, Z., Rolish, M. E., Yeo, G., Tung, V., Mawson, M., and Burge, C. B. Systematic identification and analysis of exonic splicing silencers. Cell, 119(6):831–845, 2004. doi: 10.1016/j.cell.2004.11.010.

Warner, K. D., Hajdin, C. E., and Weeks, K. M. Principles for targeting rna with drug-like small molecules. Nature Reviews Drug Discovery, 17(8):547–558, 2018. doi: 10.1038/nrd.2018.93.

Wilkinson, M. E., Charenton, C., and Nagai, K. Rna splicing by the spliceosome. Annual Review of Biochemistry, 89:359–388, 2020. doi: 10.1146/annurev-biochem-091719-064225.

Wilks, C., Gaddipati, P., Langmead, B., and Leek, J. T. Snaptron: querying splicing patterns across tens of thousands of rna-seq samples. Bioinformatics, 34(1):114–116, 2018. doi: 10.1093/bioinformatics/btx547.

Wong, M. S., Kinney, J. B., and Krainer, A. R. Quantitative activity profile and context dependence of all human 5’ splice sites. Molecular Cell, 71(6):1012–1026.e3, 2018. doi: 10.1016/j.molcel.2018.08.015.

Zhang, T. T. et al. High-throughput fluorescence polarization method for identifying ligands of lox-1. Acta Pharmacologica Sinica, 27(4):447–452, 2006. doi: 10.1111/j.1745-7254.2006.00304.x.

Zhu, M. R. et al. Development of a high-throughput fluorescence polarization assay for the discovery of ezh2-eed interaction inhibitors. Acta Pharmacologica Sinica, 39(2):302–310, 2018. doi: 10.1038/aps.2017.117.

